# FGL-1 binding to LAG-3 inhibits T cell activation via disruption of CD28 and TCR signaling

**DOI:** 10.1101/2025.08.05.668721

**Authors:** Samantha Nyovanie, Anika Hashem, Evgeny Kanshin, Nicolas Oltean, Beatrix Ueberheide, Yury Patskovsky, Michelle Krogsgaard

## Abstract

Lymphocyte-activation gene 3 (LAG-3) is known to suppress T cell receptor (TCR) signaling by disrupting CD4/CD8 coreceptor and Lck association in the absence of ligand binding. However, the impact of Fibrinogen-like protein 1 (FGL-1) binding in modulating LAG-3 mediated inhibition and its specific impact on CD28 costimulatory signaling remains unclear. This study investigates the role of LAG-3/FGL-1 interaction in modulating T cell activation. Using phosphoproteomics and confocal microscopy in LAG-3+ Jurkat cells, we demonstrated that LAG-3/FGL-1 interaction suppresses T cell activation by disrupting CD28-mediated recruitment of Lck, impairing its colocalization with CD28. Importantly, this inhibition occurs independently of CD4-bound Lck, indicating that LAG-3 ligand engagement targets both free and membrane-associated Lck pools. Consequently, this leads to suppressed phosphorylation of CD28 and TCR/CD3 signaling components, broadly attenuating downstream tyrosine phosphorylation and T cell effector functions. Our findings identify the CD28-Lck axis as a critical target of LAG-3/FGL-1 inhibitory pathway and provide a rationale for therapeutic strategies that reinvigorate CD28-Lck signaling to enhance anti-tumor immunity.

## INTRODUCTION

Effective T cell activation is initiated by the engagement of the T cell receptor (TCR) with antigenic peptide-major histocompatibility complex (pMHC) and is further strengthened by a secondary signal from costimulatory receptors such as CD28.^1–3^ The interaction between CD28 and its ligands, CD80 and CD86, is essential; in the absence of costimulatory signaling, T cells enter a state of anergy and lose their ability to respond to antigenic stimulation.^4,5^ Upon concurrent TCR and CD28 ligand engagement, the Src-family protein tyrosine kinase Lck is recruited to the immunological synapse (IS), where it phosphorylates tyrosine residues within CD28 and the immunoreceptor tyrosine-based activation motifs (ITAMs) in CD3.^3,6,7^ This phosphorylation initiates a cascade of signaling events including activation of ZAP-70 and other downstream molecules.^3,6–8^ Furthermore, CD28 co-stimulation has been shown to sustain Lck activation via autophosphorylation at Y394 and promote its recruitment and retention at IS.^9–12^ This, in turn, increases the phosphorylation of CD3 ITAMs and ZAP-70, amplifying TCR/CD3 signaling.

T cell activation is tightly regulated by co-inhibitory receptors (IRs) such as programmed death-1 (PD-1) and lymphocyte-activation gene 3 (LAG-3) to prevent overactivation and autoreactivity. Tumors often exploit these inhibitory pathways by expressing corresponding ligands, thereby dampening the anti-tumor immune response. LAG-3 interacts with several known ligands including pMHC-II, Galectin-3 (Gal-3), liver sinusoidal endothelial cell lectin (LSECtin) and fibrinogen-like protein 1 (FGL-1).^13–16^ Particularly, LAG-3 interaction with stable pMHC class II and FGL-1 has been shown to suppress T cell proliferation and cytokine production.^17,18^ MHC II is the best characterized ligand, and the structure of mouse LAG-3 in complex with pMHC II (I-Ab) demonstrates that the D1 domain of LAG-3 binds a conserved, membrane-proximal region of MHC II, spanning both the α2 and β2 subdomains.^19–21^ LAG-3 dimerization appears to restrict the spacing of pMHC II molecules at the immunological synapse, potentially attenuating T cell activation by imposing suboptimal signaling geometry.^21,22^ FGL-1, a secreted glycoprotein belonging to the fibrinogen superfamily, is predominantly expressed in hepatocytes under normal physiological conditions.^23–25^ However, FGL-1 is frequently upregulated in various solid tumors, elevated levels have been associated with poor clinical outcomes.^17^ Structure-function studies show that pMHC II and FGL-1 bind to distinct but potentially overlapping regions on LAG-3, and both interactions suppress T cell proliferation.^17,21^ Upon ligand binding, LAG-3 transmits inhibitory signals via its intracellular domain, which lacks canonical tyrosine-based inhibitory motifs (ITIMS and ITSMs) found in other IRs. Instead, the LAG-3 intracellular domain (ICD) contains several unique conserved motifs: the FxxL motif, KIEELE sequence, and a negatively charged glutamic acid–proline repeat (EP repeat), that have been shown to be essential for LAG-3-mediated T cell suppression.^26–29^

Recent studies have provided important mechanistic insights into how LAG-3 exerts its inhibitory effects. Jiang et al. demonstrated that pMHC II and FGL-1 binding triggers LAG-3 lysine-ubiquitination in the ‘KIEELE’ sequence, leading to release of its cytoplasmic tail from the membrane and activation of the LAG-3 inhibitory pathway.^30^ In parallel, Guy et al, reported that LAG-3 may exert inhibitory effects independent of ligand binding.^31^ In the absence of ligand binding, LAG-3 associates with the TCR/CD3 complex and disrupts the interaction between Lck and CD4/CD8 coreceptors via its cytoplasmic domain. Specifically, the EP repeat lowers the local pH at the IS and chelates Zn^2+^ ions from Lck-coreceptor complexes, thereby limiting Lck recruitment to the synapse.^31^ Further, Du et al., demonstrated that effective LAG-3/pMHC II inhibitory effect requires spatial proximity to the TCR, achieved through cognate pMHC II presentation and is independent of CD4 coreceptor engagement.^32^ They also identified a direct interaction between the LAG-3 intracellular domain and CD3ε via the FxxL motif, which disrupts CD3ε/Lck condensates. Despite these advances, it remains unknown whether LAG-3 directly impairs CD28 costimulatory signaling, potentially through modulation of Lck activity. Moreover, the precise mechanisms and the extent to which LAG-3 independently inhibits TCR/CD3 and CD28 signaling pathways, beyond CD4/CD8-mediated Lck recruitment, remains to be fully elucidated.

This study aims to address critical knowledge gaps by investigating the functional significance of FGL-1 binding in the LAG-3 inhibitory pathway. To dissect the molecular mechanisms underlying LAG-3/FGL-1 interactions and downstream signaling, we employed a combination of biochemical assays, phosphoproteomic analysis, and confocal microscopy in Jurkat T cells engineered for constitutive LAG-3 expression.

We first validated the biological activity of recombinant FGL-1 by demonstrating its binding to LAG-3 and its potent inhibitory effect on CD3/CD28-mediated T cell activation. Phosphoproteomic analyses revealed that LAG-3/FGL-1 interaction significantly suppresses phosphorylation of proximal signaling components downstream of TCR/CD3 and CD28, leading to a broad reduction in phosphotyrosine (pY) signaling. Complementary confocal microscopy demonstrated enhanced colocalization of LAG-3 with CD28 upon T cell activation, indicating spatial proximity and a direct molecular interaction. Notably, the colocalization between CD28 and active Lck (pY394^9^) was significantly reduced upon FGL-1 stimulation. These findings suggest that LAG-3/FGL-1 engagement disrupts Lck pY394 association and CD28 phosphorylation. Given the critical role of CD28 costimulatory signaling in recruiting and activating Lck at the IS,^10,11^ our data highlight a multifaceted inhibitory role of LAG-3 that targets key nodes in both TCR/CD3 and CD28 signaling pathways to suppress T cell activation.

In the broader context, our findings contribute to the rapidly evolving landscape of immune checkpoint biology. By elucidating the molecular interactions through which LAG-3 modulates TCR signaling, we provide critical insights that may inform the development of next-generation therapeutic strategies aimed at enhancing anti-tumor immunity while minimizing immune-related adverse events. Moreover, this work underscores how multiple inhibitory receptors, such as LAG-3 and PD-1, may cooperate synergistically to regulate T cell activation and exhaustion by simultaneously targeting both antigenic and costimulatory pathways. Targeting the LAG-3/FGL-1 axis thus offers a promising strategy to improve therapeutic outcomes and expand the scope of effective immunotherapies.

## RESULTS

### Recombinant FGL-1 fibrinogen domain binds LAG-3 and suppresses T cell activation

To investigate the functional impact of LAG-3/FGL-1 interaction on T cell activation, we generated the recombinant Fibrinogen-like domain (FD) of FGL-1, which has previously been shown to bind to LAG-3.^17^ Given that full-length FGL-1 forms higher-order oligomers via its coiled-coil domain ^25,33,34^ we expressed FGL-1 FD with a C-terminal Avi-Tag to enable site-specific biotinylation,^35^ and subsequent multimerization via streptavidin. We purified FGL-1 FD by size exclusion chromatography (SEC), which resolved distinct peaks corresponding to dimeric (61.5 kDa) and monomeric (30.75 kDa) species (**Fig. S1 A**). Further purification confirmed protein purity and structural integrity (**Fig. S1 B, C**). The monomeric fraction exhibited doublet bands, consistent with variable cysteine oxidation states. To stabilize the FGL-1 FD monomer, we included 2 mM CaCl_2_ in the refolding and purification buffers, resulting in a monodispersed peak on SEC (**Fig. S1 A** and **Fig. S1 D**). Biotinylation efficiency was confirmed by gel-shift assay using SDS-PAGE, which showed a protein-streptavidin complex (**Fig. S1 E**), indicating successful and functional biotinylation.

To characterize the structural differences, we determined the structure of FGL-1 FD dimer at a resolution of 1.74 Å (**Fig. S1 F, Table S1**). The dimer adopts a conformation stabilized by an interchain disulfide bond, which likely enhances LAG-3 interaction through avidity effects. Comparative analysis with the previously published FGL-1 FD monomer structure (PDB: 7TZ2)^36^ revealed overall structural similarity, with notable differences localized to the dimer interface and the calcium-binding site (**Fig. S1G**). Interestingly the FGL-1 FD loss of function (LOF) mutation: Y244A/E245A (corresponding to Y176/E177 in FGL-1 FD dimer), identified by Ming et., al^36^, is located near the dimer interface, yet remains solvent exposed (**Fig. S1 G**). Using Bio-Layer Interferometry (BLI), we assessed binding affinities of LAG-3 to monomeric and dimeric FGL-1 FD. The dimer exhibited substantially higher affinity (K_D_ = 16.8 nM) compared to FGL-1 FD monomer (K_D_ = 0.74 µM) (**Fig. S1 H, I**), demonstrating that FGL-1 FD dimerization dramatically enhances LAG-3 binding. Notably, this increased affinity occurs in the absence of calcium, indicating that dimerization/multimerization itself is a key determinant in optimizing LAG-3/FGL-1 interaction.

To validate the specific interaction between LAG-3 and FGL-1, we performed tetramer binding assays using Jurkat T cell line stably expressing LAG-3. Fluorescently labeled FGL-1 FD::streptavidin Phycoerythrin (SA-PE) tetramers were titrated onto Jurkat cells, and LAG-3/FGL-1 FD binding was quantified by flow cytometry. The tetramer assay demonstrated concentration-dependent and specific binding of FGL-1 to LAG-3+ Jurkat cells (**Fig. 1 A**), with significantly stronger binding compared to wild-type (WT) Jurkat cells, which express low levels of LAG-3 (**Fig. 1 B**). Similar binding patterns were observed with FGL-1 FD dimer on LAG-3+ Jurkat cells (**Fig. S2A**), indicating that multimerization via streptavidin effectively may recapitulate the physiological higher-order oligomers of FGL-1. Our results confirm that FGL-1 FD binds LAG-3 with high affinity, thereby validating the functional interaction between LAG-3 and FGL-1.

**Figure 1.**
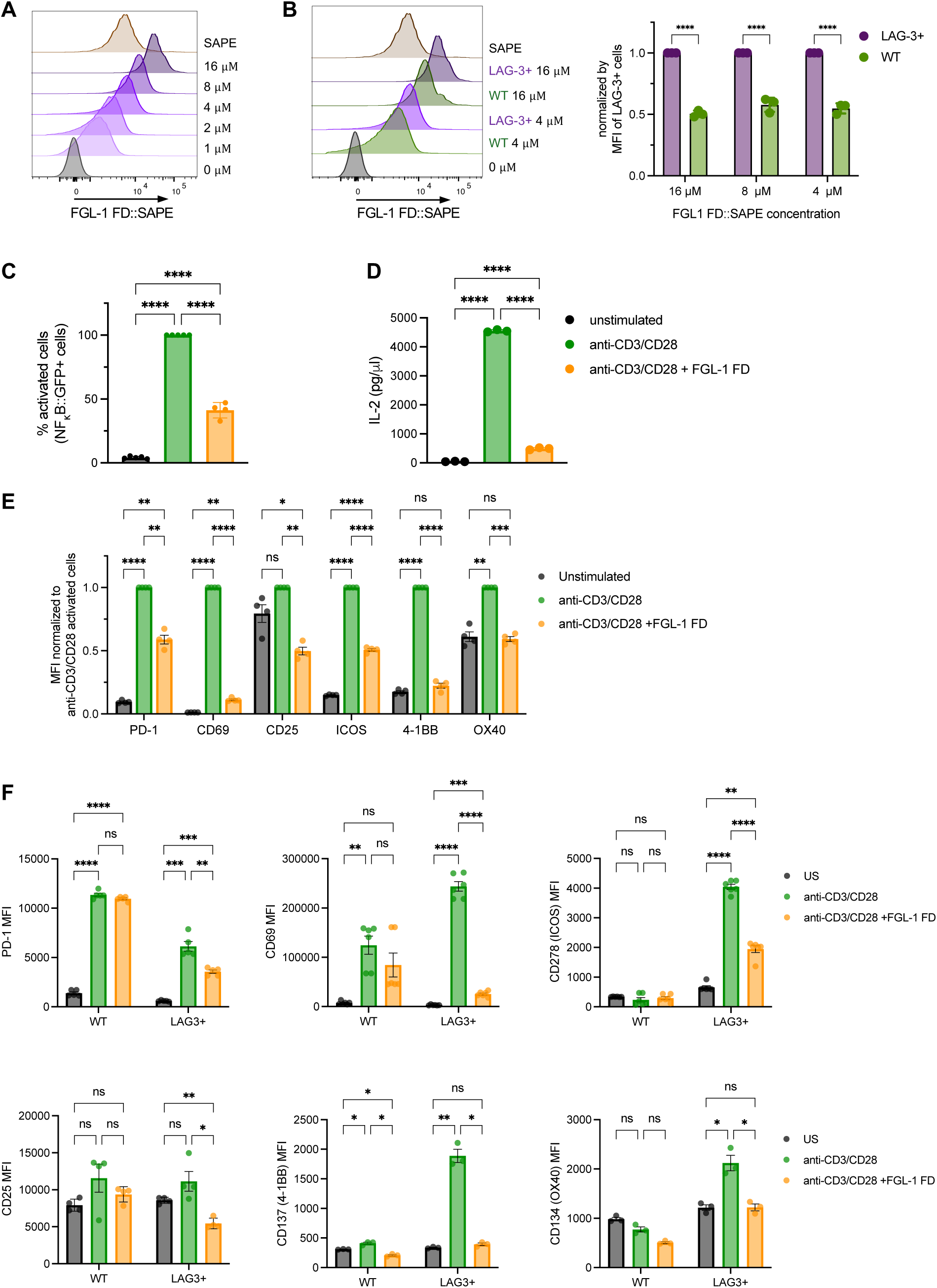
Recombinant FGL-1 fibrinogen domain binds LAG-3 and suppresses T cell activation. **(A)** Flow cytometry analysis of tetrameric FGL-1 FD::streptavidin Phycoerythrin (SA-PE) binding to LAG-3+ Jurkat cells across a concentration range (1 – 16 µM). **(B)** Comparison of FGL-1 FD tetramer binding to LAG-3+ Jurkat cells versus wild-type (WT) Jurkat cells (LAG-3 low) (n = 3). Data are presented as median fluorescent intensity (MFI), normalized to LAG-3+ Jurkat cells MFI at each concentration. Error bars represent standard deviation (SD). **(C)** NF-κB::eGFP induction and **(D)** IL-2 production in LAG-3+ Jurkat T cells stimulated with anti-CD3/CD28-coated beads +/− FGL-1 FD or mouse IgG1 isotype control (mIgG1) for 24 hours (n = 4, n = 3, respectively). NF-κB::eGFP induction data is shown as percent MFI normalized to anti-CD3/CD28 stimulation (set to 100%). Error bars represent standard deviation (SD). **(E)** Expression of T cell activation markers: PD-1, CD69, CD25, ICOS, 4-1BB and OX40 in LAG-3+ Jurkat T cells, either unstimulated or stimulated with anti-CD3/CD28 +/− FGL-1 FD (n = 4). Data are shown as MFI ± SEM, normalized to the anti-CD3/CD28 condition. **(F)** Control expression of activation markers in WT (LAG-3 low) Jurkat T cells under identical conditions (n = 4). Data represent MFI ± SEM from triplicate wells across two independent experiments. Statistical significance was performed using one or two-way ANOVA, followed by Tukey or Šídák’s post hoc tests, respectively. **** p < 0.0001, *** p < 0.001, ** p < 0.01, * p < 0.05, ns = not significant p > 0.05.

Next, we assessed the functional impact of FGL-1 on T cell activation using LAG-3+ Jurkat NF-κB::eGFP reporter cells stimulated with anti-CD3 and anti-CD28 antibodies in the presence of FGL-1 or mouse IgG1 (mIgG1) isotype control coated beads (**Fig. S2 B**). NF-κB::eGFP expression was significantly reduced in cells stimulated with FGL-1 FD, showing a mean decrease of 59% compared to stimulation without FGL-1 FD (**Fig. 1 C, Fig. S2 C, D**). In addition, FGL-1 FD markedly suppressed IL-2 production (**Fig. 1 D, Fig. S2 E**) and expression of activation-induced surface markers including PD-1, CD69, CD25, ICOS, 4-1BB and OX40, as measured by flow cytometry (**Fig. 1 E**). This inhibitory effect was specific to the LAG-3/FGL-1 FD interaction, as WT Jurkat cells showed no significant effect of FGL-1 on activation markers expression (**Fig. 1 F**).

To further confirm this interaction and its functional consequences, we generated FGL-1 FD LOF mutant, Y244A/E245A. This LOF mutant exhibited markedly reduced tetramer binding (**Fig. S3 A**) and significantly reduced T cell suppression, as evidenced by higher NF-κB::eGFP induction and IL-2 production compared to FGL-1 FD (wild-type) (**Fig. S3 B, C**). These findings further underscore the specificity of the LAG-3/FGL-1 interaction. Collectively, these results demonstrate that FGL-1 FD binds LAG-3 to mediate suppression of CD3/CD28-driven T cell activation, contrasting with previous reports suggesting that LAG-3 mediated inhibition can occur independently of ligand engagement.^31^

### LAG-3/FGL-1 FD interaction suppresses global T cell tyrosine phosphorylation

Having established that the LAG-3/FGL-1 FD interaction suppresses T cell activation, we next hypothesized that FGL-1 engagement initiates LAG-3-mediated inhibitory signaling, thereby amplifying its suppressive effect. To test this, we examined the impact of LAG-3/FGL-1 FD interaction on proximal TCR/CD3 signaling (Lck and ZAP-70 phosphorylation) and downstream MAPK signaling (ERK-1/2 phosphorylation) in LAG-3+ Jurkat cells stimulated for 5 – 15 minutes. Western blot analysis revealed FGL-1 FD presence resulted in a significant decrease in the phosphorylation of Lck at Y394, ZAP-70 at Y319, and ERK-1/2 at T202/Y204 (**Fig. S4**), indicating. broad attenuation of TCR/CD3 signaling. Notably, these changes were most pronounced within the first 5 minutes of stimulation, suggesting that LAG-3/FGL-1-mediated inhibition occurs rapidly and targets early T cell activation events. Since both FGL-1 FD monomer and dimer elicited comparable suppression of T cell activation and proximal signaling, subsequent experiments were conducted using monomeric form exclusively to reduce redundancy. Hereafter, “FGL-1 FD” refers to the monomeric species.

Given these findings, we employed a phosphoproteomic approach to gain deeper insights into the molecular mechanism underlying the LAG-3/FGL-1-mediated signaling inhibition. Phosphopeptide detection is inherently challenging due to their low abundance, with phosphotyrosine (pY) sites being particularly rare and difficult to identify.^37^ To overcome this limitation, we optimized pY peptide enrichment using an engineered Src Homology 2 superbinder domain (SH2 SB), which has significantly higher affinity and broader specificity for pY-containing peptides compared to wild-type SH2 domains.^38–40^ Originally developed by Kaneko et al., this SH2 SB incorporates three-point mutations (Thr183Val/Ser188Ala/Lys206Leu) in the pY-binding pocket, improving its binding performance. To increase recombinant SH2 stability and reduce possible oxidation-induced heterogeneity, we introduced two additional cysteine-to-serine substitutions (Cys241Ser/Cys248Ser) to eliminate reactive thiol groups.

The SH2 SB mutant was expressed and purified (**Fig. S5 A, B**). We confirmed its structural integrity and pY-binding capacity using high-resolution X-ray crystallography and binding assays. The crystal structures of ligand-free SH2 SB and SH2 SB bound to O-Phospho-L-tyrosine were resolved at 1.50 Å and 1.61 Å resolution, respectively, and exhibited remarkable conformational similarity, with a root-mean-square deviation (RMSD) of 0.137 (**Fig. S5 C – E, Table S2**). Detailed analysis of the pY-binding pocket revealed that the pY phosphate group is stabilized by a series of hydrogen bonds with SH2 SB residues Arg158, Arg178, Ser180, and Thr182 (**Fig. S5 F**). Structural superposition with the SH2 SB reported by Kaneko et al., yielded RSMDs of 0.204 Å (unbound) SH2 SB and 0.229 Å (pY-bound), confirming high structural conservation. Importantly, the Cys241Ser/Cys248Ser mutations did not affect the structure or the pY binding ability of the SH2 SB (**Fig. S5 G-H**).

To evaluate the binding capacity and sensitivity of the mutant SH2 SB for phosphotyrosine (pY) peptides relevant to T cell signaling, we first generated tryptic peptide sequences of key TCR signaling proteins: ZAP-70, Lck, ERK1, and ERK2, using PEPTIDEMASS^41^ Based on these sequences, we designed four synthetic pY peptides incorporating critical TCR associated pY sites (**Table S3**). We performed enrichment assays by incubating the resin with immobilized SH2 SB with titrated amounts of synthetic pY peptides spiked into a HeLa cell peptide digest standard to mimic a complex biological mixture. The mutant SH2 SB efficiently captured these pY peptides across a wide concentration range, demonstrating a clear correlation between peptide concentration and signal intensity (**Fig. S5 H**). Importantly, this high-affinity binding was maintained even in the presence of complex peptides mixtures, underscoring the robustness and specificity of the engineered SH2 SB for capturing low-abundance pY peptides. Next, we compared pY peptide enrichment using SH2 SB only (pY SB) versus a combined Ti^4+^-IMAC and SH2 SB (pY IMAC-SB) from H_2_O_2_ treated Jurkat lysates. Dong et al. previously reported that including the Ti^4+^-IMAC enrichment step improved the number pY peptides identified by 62 % and increased specificity to 90%.^39^ However, our results showed that the pY SB method identified more pY peptides (6,855) with higher specificity (49%) than the combined pY IMAC-SB method (3,906 peptides, 11% specificity) (**Fig. S5 I, Table S4**). Ti^4+^-IMAC did not effectively remove all non-phosphorylated peptides and caused loss of lower abundance of pY peptides. Moreover, the majority of the pY peptides identified by the pY IMAC-SB method (92%, 3586 peptides) were also detected using the SH2 SB only enrichment method (**Fig. S5 J**). Overall, SH2 SB-only enrichment offers a simpler, more effective, and more comprehensive approach for enrichment of pY peptides. In summary, our data confirm the structural integrity and functionality of the mutant SH2 SB, supporting its utility in phosphoproteomics enrichment experiments.

We next applied peptide digests from unstimulated and stimulated LAG-3+ Jurkat cells to SH2 SB-immobilized resin, followed by mass spectrometric analysis of the enriched pY peptides. We identified 1,566 unique pY peptides across 660 proteins. Notably, FGL-1 FD-treated cells exhibited reduction in both the number and intensity of recovered pY peptides (180 – 487 peptides) compared to cells stimulated without FGL-1 FD (314 – 630 peptides) from equivalent protein inputs (**Fig. 2 A, B**). Only 31% of pY peptides identified in anti-CD3/anti-CD28 stimulated cells (261 peptides) were also detected in cells co-stimulated with FGL-1 FD (**Fig. 2 C**). Among these, no statistically significant differences in phosphorylation were observed across unstimulated and stimulated conditions (± FGL-1 FD) at a false discovery rate (FDR) < 5% (**Fig. 2 D – F**). Despite the absence of significant site-specific phosphorylation changes, the consistent reduction in both the number and abundance of pY peptides following FGL-1 FD stimulation suggests that the LAG-3/FGL-1 FD broadly suppresses global tyrosine phosphorylation along the TCR signaling pathway. These results provide additional evidence that LAG-3 ligation by FGL-1 FD contributes to the observed inhibition of T cell activation by attenuating early tyrosine phosphorylation signaling events.

**Figure 2.**
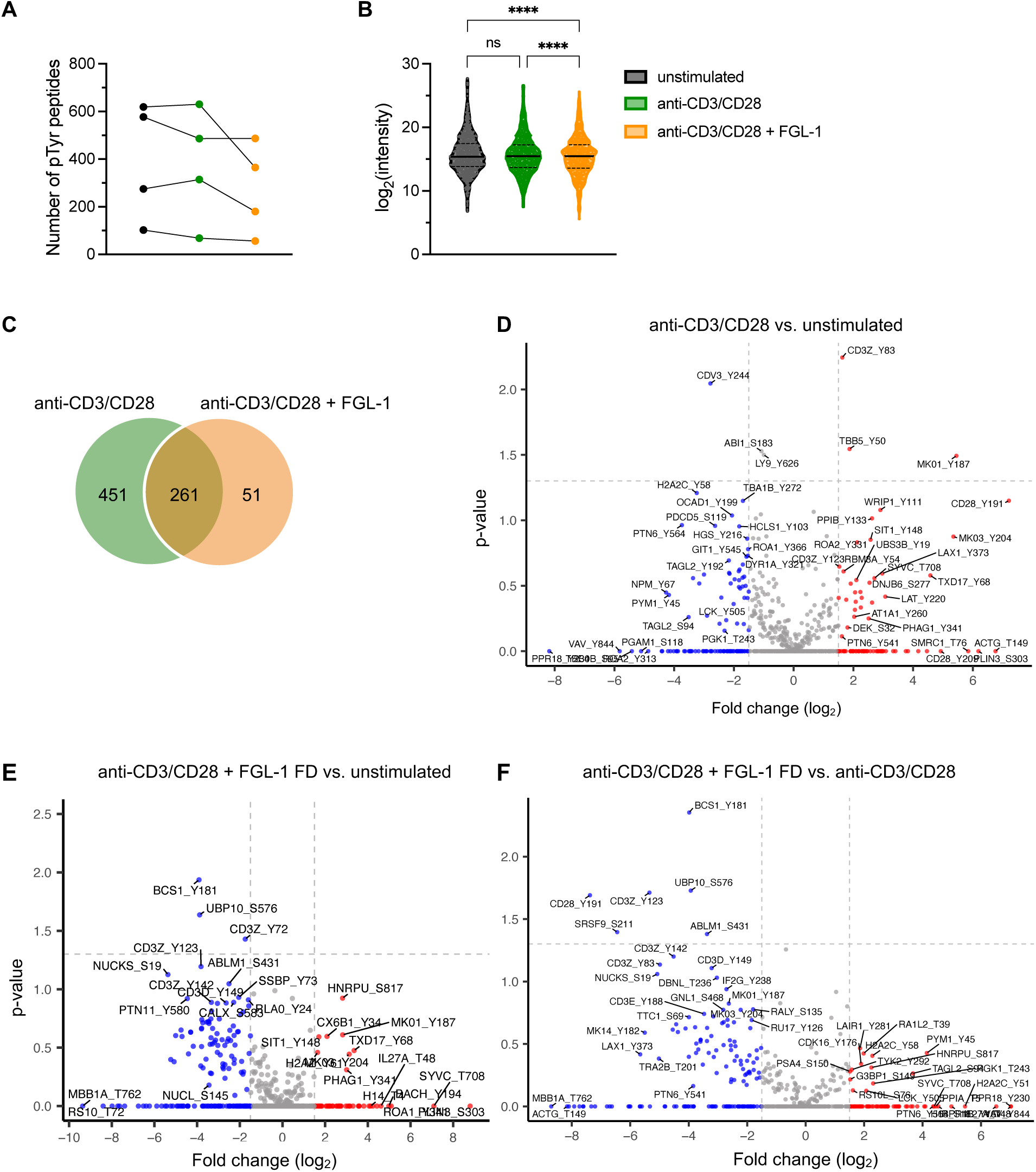
LAG-3/FGL-1 FD interaction globally suppresses T cell tyrosine phosphorylation. **(A, B)** Label-free LC–MS/MS analysis of phosphotyrosine (pY) peptides in LAG-3+ Jurkat cells, either unstimulated or stimulated for 5 minutes with anti-CD3/CD28 +/− FGL-1 FD (n = 4), **(A)** number of unique pY peptides identified per condition **(B)** Log_2_ intensity distribution of detected pY peptides. Statistical analysis performed using two-way ANOVA with Tukey’s post hoc tests. Solid black line = median; dashed lines = interquartile range. **** p < 0.0001, ns = not significant (p > 0.05) **(C)** Venn diagram showing overlap of pY peptides detected in cells stimulated with anti-CD3/anti-CD28 versus anti-CD3/anti-CD28 + FGL-1 FD from one representative experiment. **(D – F)** Volcano plots illustrating differential log_2_ fold changes (log_2_FC) in pY intensity differences, across the three conditions: **(D)** anti-CD3/CD28 vs. unstimulated; **(E)** anti-CD3/CD28 + FGL-1 FD vs. unstimulated; **(F)** anti-CD3/CD28 + FGL-1 FD vs. anti-CD3/CD28. Red = significantly upregulated phosphosites (log_2_FC ≥ 1.5); blue = significantly downregulated phosphosites, (log_2_FC ≤ 1.5); grey = unchanged phosphosites (log_2_FC) < 1.5).

### Quantitative phosphoproteomics reveals LAG-3/FGL-1-mediated suppression of proximal TCR signaling

To further investigate the phosphoproteomic changes induced by LAG-3/FGL-1 engagement, we analyzed the log_2_ fold change (log_2_FC) values for all identified phosphorylation sites across the three stimulation conditions (**Fig. 2 D – F**). As an initial step, proteins exhibiting pY site log_2_FC values of ±1.5 were selected for functional enrichment analysis using the Database for Annotation, Visualization and Integrated Discovery (DAVID) Bioinformatics Resources.^42,43^ This fold-change threshold was applied solely to identify perturbed biological pathways and does not imply definitive regulation of individual phosphosites, especially since no pY sites met an FDR <5%. Pathway analysis using the Kyoto Encyclopedia of Genes and Genomes (KEGG) revealed that mRNA processing and TCR signaling were the most significantly enriched pathways following stimulation in the presence of FGL-1 FD, compared to both T cell stimulation in the absence of FGL-1 FD and unstimulated controls (**Fig. 3 A**).

**Figure 3.**
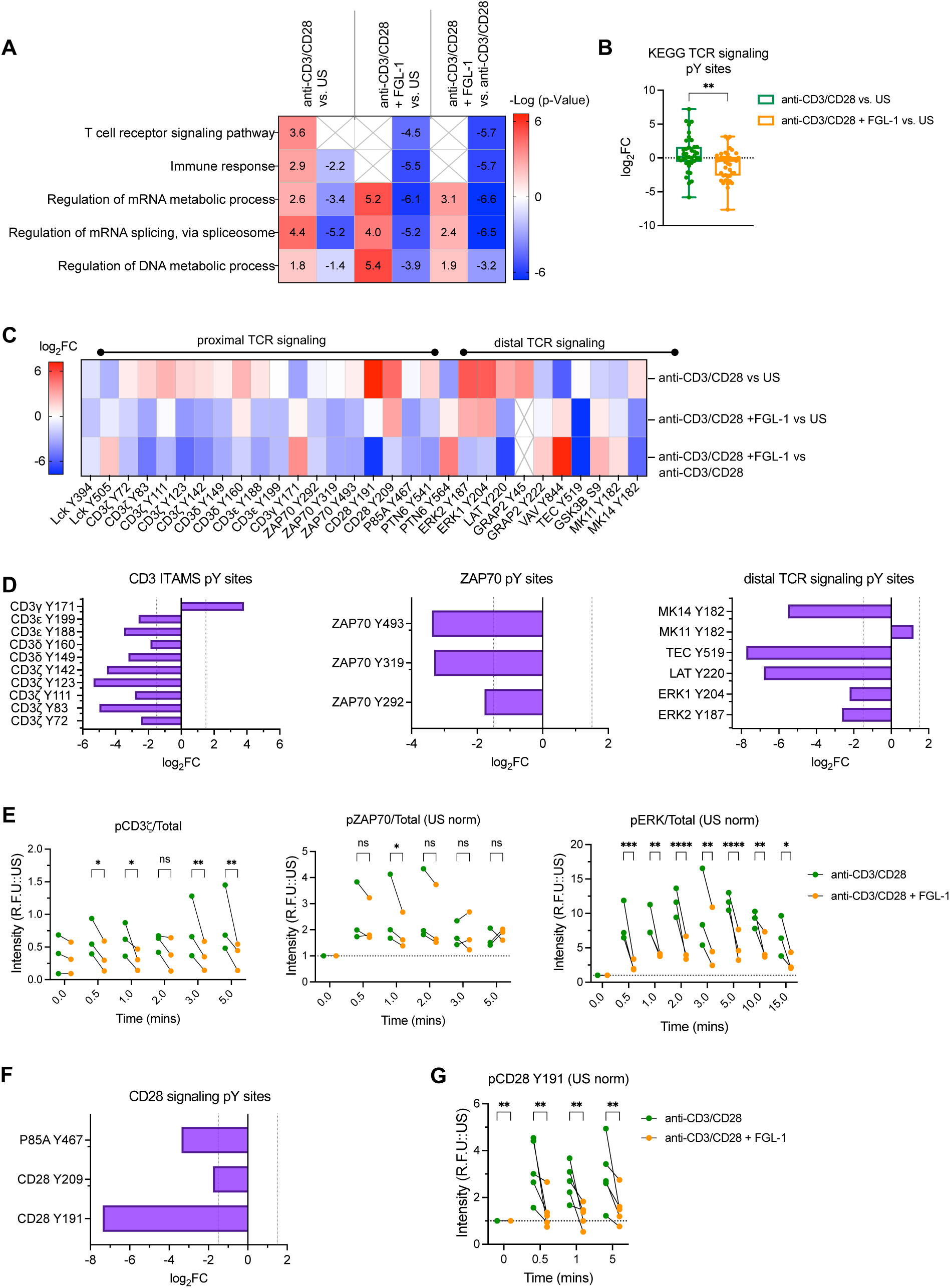
LAG-3/FGL-1 interaction inhibits TCR and CD28 signaling pathways. (**A**) Functional enrichment analysis of proteins with upregulated pY sites (log_2_FC ≥ 1.5, red), and downregulated pY (log_2_FC ≤ 1.5; blue) in response to anti-CD3/CD28 and anti-CD3/CD28 + FGL-1 FD stimulation, relative to unstimulated cells and stimulation without FGL-1 FD. Color intensity indicates –Log_10_ (p-value) of pathway enrichment; X denotes non-enriched pathways. (**B**) Log_2_ fold change (log_2_FC) analysis of TCR signaling-associated pY peptides following anti-CD3/CD28 and anti-CD3/CD28 + FGL-1 FD stimulation compared to unstimulated cells. (**C**) Heatmap of log_2_FC phosphorylation differences in TCR signaling-related pY peptides across three conditions: anti-CD3/CD28 vs. unstimulated; anti-CD3/CD28 + FGL-1 FD vs. unstimulated; anti-CD3/CD28 + FGL-1 FD vs. anti-CD3/CD28. (**D**) Bar graphs showing log_2_FC values of TCR signaling pY sites in CD3 ITAMS, ZAP-70, LAT, ERK1/2, TEC and distal molecules upon FGL-1 FD stimulation relative to stimulation without FGL-1 FD. (**E**) Time-course western blot analysis of phosphorylation at CD3ζ Y142, ZAP70 Y319, and ERK 1/2 Y204/Y187 in LAG-3+ Jurkat cells unstimulated, stimulated with anti-CD3/CD28 +/− FGL-1 FD. Quantification represents phospho/total protein signal intensity, normalized to unstimulated (US) values (n = 3). (**F**) Bar graphs showing log_2_FC values of CD28 signaling pY sites: CD28 pY191, pY209, and PI3K P85A pY467, in response to FGL-1 FD stimulation relative to stimulation without FGL-1 FD. (**G**) Western blot analysis of CD28 Y191 phosphorylation in LAG-3+ cell lysates: unstimulated, stimulated with anti-CD3/CD28 +/− FGL-1 FD. Band intensities were quantified and normalized to total CD28 and unstimulated control (n = 5). Statistical significance was determined using a two-way ANOVA with Šídák’s post hoc tests. **** p < 0.0001, *** p < 0.001, ** p < 0.01, * p < 0.05, ns = not significant (p > 0.05).

These findings indicate that FGL-1 FD engagement broadly impacts both post-transcriptional regulation and canonical T cell signaling cascades. Consistent with this, we observed a marked reduction in phosphorylation at key TCR signaling protein pY sites after T cell activation in the presence of FGL-1 FD as compared to stimulation without FGL-1 FD (**Fig. 3 B**). Specifically, proximal TCR signaling components, including CD3 ITAMS and ZAP-70 pY sites, exhibited decreased phosphorylation levels (**Fig. 3 C**). Among the five detected CD3ζ pY sites, three (CD3ζ Y83, Y123, and Y142) showed significantly reduced phosphorylation upon LAG-3/FGL-1 FD stimulation (**Fig. S6 A**). Similarly, phosphorylation of CD3δ, CD3ε and ZAP-70 was also suppressed in the presence of FGL-1 FD (**Fig. 3 D**). This attenuation of proximal TCR signaling impaired propagation of TCR/CD3 signals, as evidenced by reduced phosphorylation of downstream signaling effectors including LAT, ERK1/2, TEC, MK11 and MK14 pY sites (**Fig. 3D**). Collectively, these results demonstrate that LAG-3/FGL-1 FD interaction disrupts TCR signaling at both early and downstream nodes, contributing to its overall inhibitory effect on T cell activation.

To quantitatively assess how LAG-3/FGL-1 interaction downregulates TCR signaling and further corroborate the phosphoproteomics results, we performed a time-course analysis of phosphorylation at key signaling nodes–CD3ζ, ZAP70 and ERK–in stimulated LAG-3+ Jurkat cells (+/− FGL-1 FD) using Western blotting (**Fig. S6 B – E**). Phosphorylation of CD3ζ at Y142, ZAP70 Y319, and ERK 1/2 Y204/Y187 was markedly reduced within 1 minute of stimulation in the presence of FGL-1-FD (**Fig. 3 E**). Notably, ERK 1/2 phosphorylation showed a significant and sustained decrease beginning as early as 30 seconds and persisting through 15 minutes in FGL-1 FD-treated cells. These findings are fully consistent with our phosphoproteomic data and further support the notion that the LAG-3/FGL-1 interaction primarily inhibits TCR proximal signaling, leading to suppression of downstream tyrosine phosphorylation.

### LAG-3/FGL-1 interaction suppresses CD28 phosphorylation and costimulatory signaling

Previous studies have shown that CD28 costimulation enhances ERK phosphorylation indirectly by amplifying CD3ζ –ZAP-70 signaling.^11,44^ Given our observations of a sustained decrease in CD3ζ and ERK phosphorylation in FGL-1 FD-stimulated cells, we examined the effect of LAG-3/FGL-1 interaction on CD28-mediated signaling using the phosphoproteomic method. FGL-1 FD-treated samples showed a reduced CD28 phosphorylation as compared to those stimulated without FGL-1 FD (**Fig. S7 A**). Among these, phosphorylation at CD28 Y191 exhibited the most pronounced suppression, with a log_2_FC of –7.37 (**Fig. 3 F**). We also observed decreased phosphorylation at additional downstream effectors of CD28 signaling. Notably, the PI3K regulatory subunit p85α at Y467 (log_2_FC –3.3) and TEC kinase at Y519 (log_2_FC –7.7) showed reduced phosphorylation in the presence of FGL-1 FD (**Fig. 3 F**). These results suggest that the LAG-3/FGL-1 interaction not only inhibits TCR/CD3 signaling but also significantly impairs CD28 phosphorylation and its associated costimulatory signaling pathway.

To validate our phosphoproteomic findings, we assessed phosphorylation status of CD28 at Y191 in LAG-3+ cells by Western blotting (**Fig. S7 B, C**). Upon T cell activation in the presence of FGL-1 FD, we observed a significant decrease in CD28 Y191 phosphorylation as early as 30 seconds post-stimulation (**Fig. 3 G**). Additionally, we noted overlapping phosphorylation profiles among CD28, CD3ζ and ZAP70 (**Fig. 3 E**), with inhibition detected within 30 seconds to 1 minute. These findings suggest that LAG-3/FGL-1 interaction concurrently suppresses both CD28 costimulatory and TCR/CD3 signaling pathways.

### LAG-3/FGL-1 interaction reduces phosphorylation of TCR and CD28 signaling molecules in primary T cells

To further validate the inhibitory role of LAG-3/FGL-1, we extended our analysis to primary T cells. Peripheral blood mononuclear cells (PBMCs) from healthy donors (donors) were stimulated with anti-CD3/CD28 antibodies, IL-2, IL-7 and IL-15 for 48 hours, followed by IL-2 rest to induce LAG-3 expression. Flow cytometric analysis of CD3+ T cells from 12 donors revealed moderate LAG-3 expression, with a median of 52% (**Fig. S8, S9**). These cells were subsequently restimulated in the presence and absence of FGL-1 FD for 5 minutes, and phosphorylation changes were analyzed in LAG-3+CD3+ T cells by flow cytometry (**Fig. S8**). Notably, FGL-1 FD treatment led to reduced phosphorylation of Lck Y394, ZAP-70 Y319. In addition, we observed a more significant decrease in phosphorylation of ERK 1/2 (T202/Y204), and PDK1 (S241), AKT (S473), key effectors downstream of CD28 signaling^2,45^ (**Fig. 4**). These results corroborate our findings in LAG-3+ Jurkat cells and further support the conclusion that the LAG-3/FGL-1 interaction attenuates T cell activation by inhibiting early events in both TCR/CD3 and CD28 signaling pathways.

**Figure 4.**
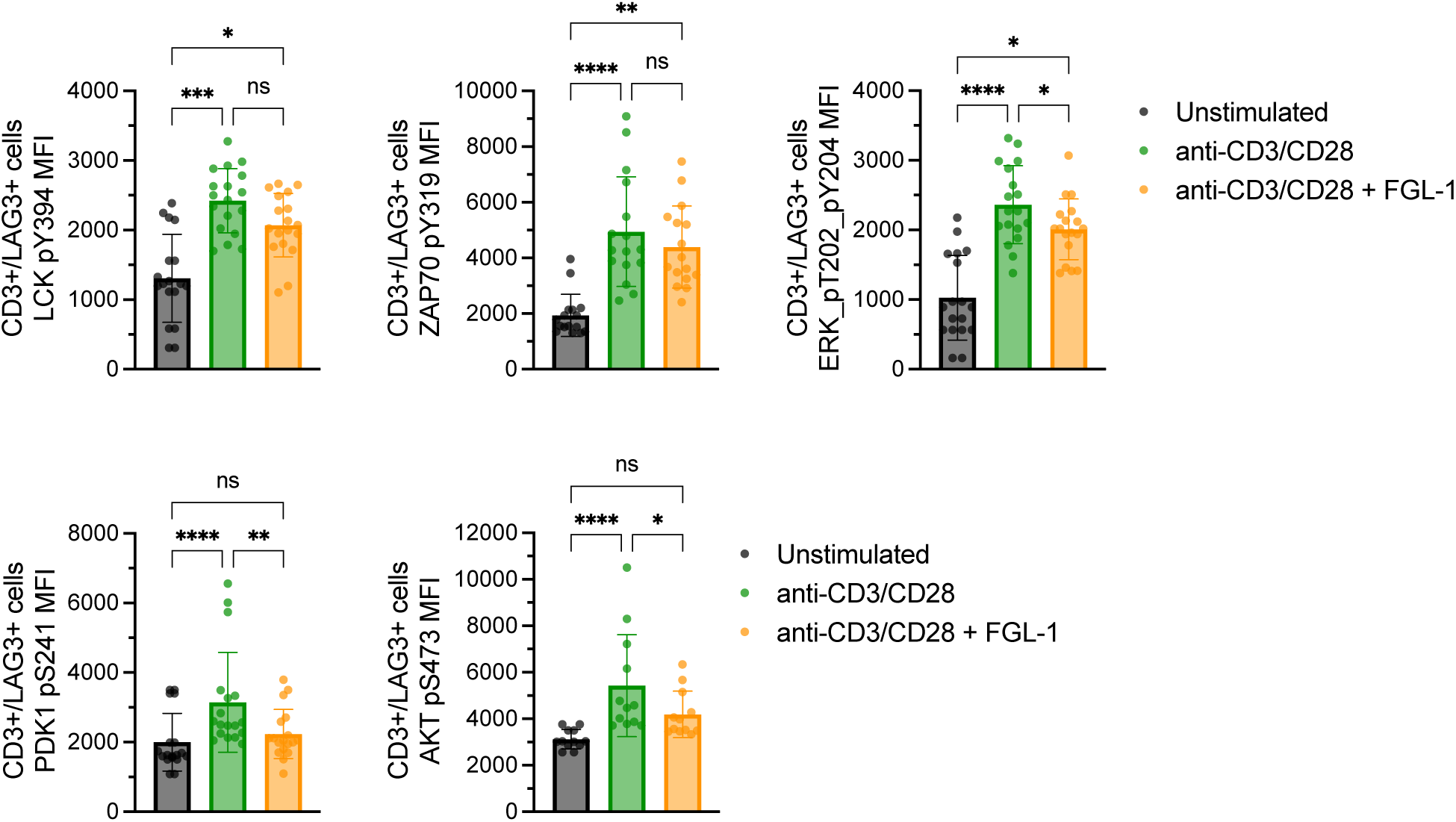
LAG-3/FGL-1 interaction reduces phosphorylation of TCR and CD28 signaling molecules in primary T cells. Phospho-flow cytometry analysis of Lck pY394, ZAP70 pY319, ERK 1/2 pT202/Y204, PDK-1 pS241, and AKT S473 in CD3+ LAG-3+ T cells from healthy donors (n ≥ 8, 2 independent experiments), under unstimulated conditions and stimulated with or without FGL-1 FD. Statistical analysis was performed using one-way ANOVA followed by Friedman’s post hoc tests. **** p < 0.0001, *** p < 0.001, ** p < 0.01, * p < 0.05, ns = not significant (p > 0.05).

### LAG-3 colocalizes with CD28 in activated Jurkat T cells

Our results indicate that LAG-3/FGL-1 interaction suppresses CD28 costimulatory signaling on a similar timescale as TCR/CD3 signaling. Given the established proximity and constitutive association of LAG-3 with the TCR/CD3 complex,^31,46^ we hypothesized that LAG-3 may similarly inhibit CD28 signaling by interfering with CD28–Lck recruitment and phosphorylation. To test this, we employed confocal microscopy to visualize the localization of TCR/CD3 and CD28 relative to LAG-3 in both unstimulated and anti-CD3/CD28-stimulated LAG-3+ Jurkat cells. Our analysis revealed that CD3 colocalized with LAG-3 to a similar extent in both unstimulated and stimulated cells, with median overlaps of 70% and 65%, respectively (**Fig. 5 A, C**). This finding aligns with previous reports demonstrating a stable association between LAG-3 and TCR/CD3 complex.^31,46^ In contrast, colocalization between CD28 and LAG-3 significantly increased upon T cell activation, with a MOC of 62% compared to 39% in unstimulated cells (**Fig. 5B, C**). Collectively, these results suggest that T cell activation promotes increased proximity between CD28 and LAG-3, potentially facilitating LAG-3 mediated inhibition of CD28 signaling.

**Figure 5.**
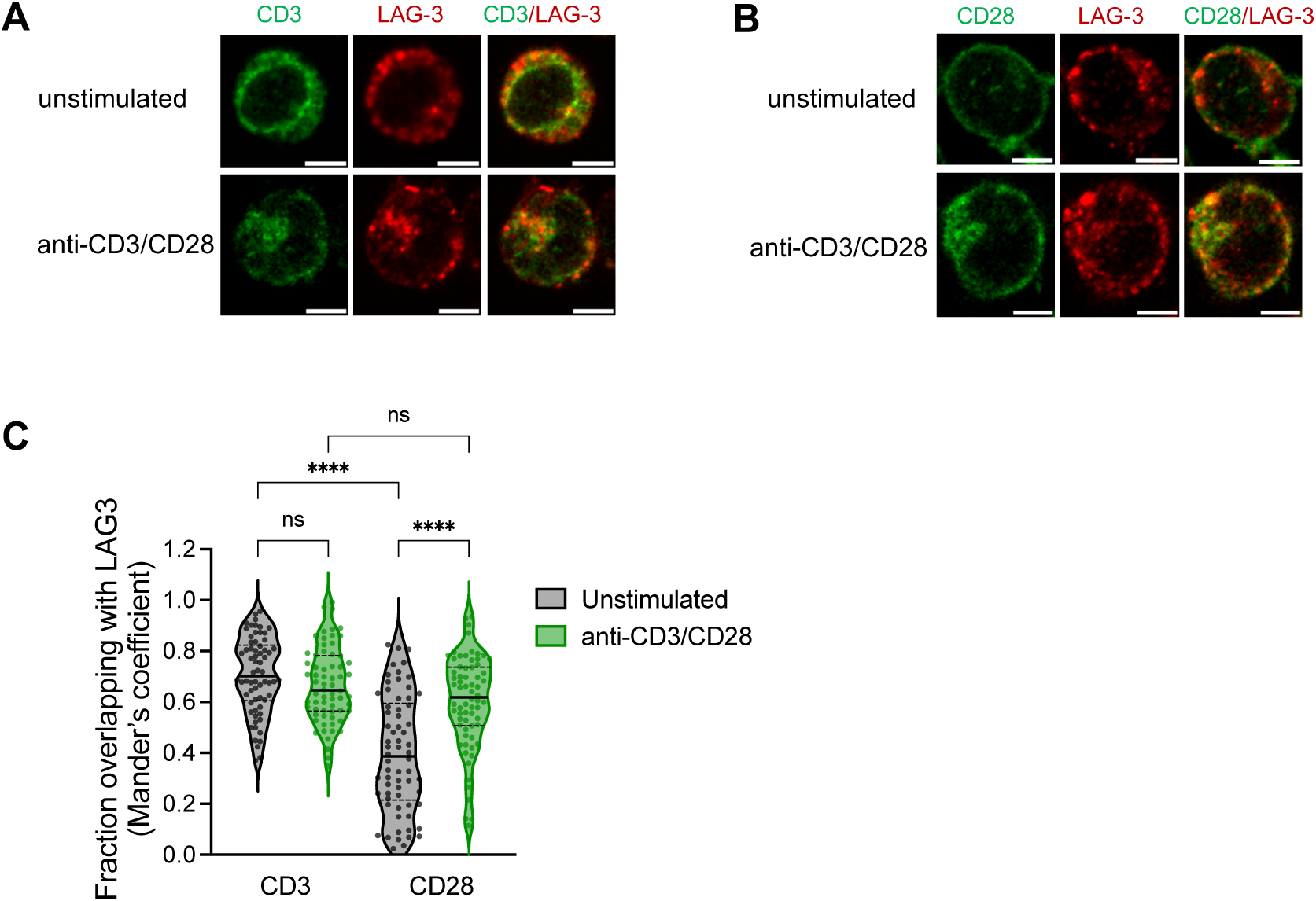
LAG-3 colocalizes with CD28 in activated Jurkat T Cells. (**A**) Representative fixed-cell confocal microscopy images showing colocalization of CD3 (green) and LAG-3 (red) in unstimulated and anti-CD3/CD28-stimulated LAG-3+ Jurkat cells. Scale bar: 5 µm. **(B)** Representative confocal images depicting colocalization of CD28 (green) and LAG-3 (red) under the same conditions: unstimulated and anti-CD3/CD28. Scale bar: 5 µm. **(C)** Violin plots of Manders’ Overlap Coefficient (MOC) quantifying colocalization of CD3 and CD28 with LAG-3 in unstimulated and stimulated cells (n = 66). Median values are indicated by solid black lines; interquartile ranges by dashed black lines. Statistical significance was assessed using Kruskal-Wallis ANOVA, followed by Dunnett’s post hoc tests. **** p < 0.0001, *** p < 0.001, ** p < 0.01, * p < 0.05, ns = not significant (p > 0.05).

### LAG-3/FGL-1 interaction downregulates CD28 and active Lck (pY394) colocalization in activated Jurkat and primary T cells

We next investigated whether the proximity of LAG-3 to CD28 influences CD28–Lck binding, and whether this effect is enhanced by ligand engagement. Specifically, we examined colocalization of CD28 with active Lck phosphorylated at Y394 (Lck pY394), which is known to specifically bind and phosphorylate CD28 upon co-stimulation.^10,11,47^ Notably, in both unstimulated and stimulated LAG-3+ cells, CD28 and Lck pY394 exhibited similar colocalization values, as measured by both MOC (median: 0.44 vs. 0.43) and PCC (median: 0.65 vs. 0.69) coefficients (**Fig 6 A, B**). However, stimulation the presence of FGL-1 FD led to a significant decrease in colocalization (MOC: 0.30, PCC: 0.61) (**Fig 6 A, B**), suggesting that the LAG-3/FGL-1 interaction disrupts CD28-Lck pY394 association.

**Figure 6.**
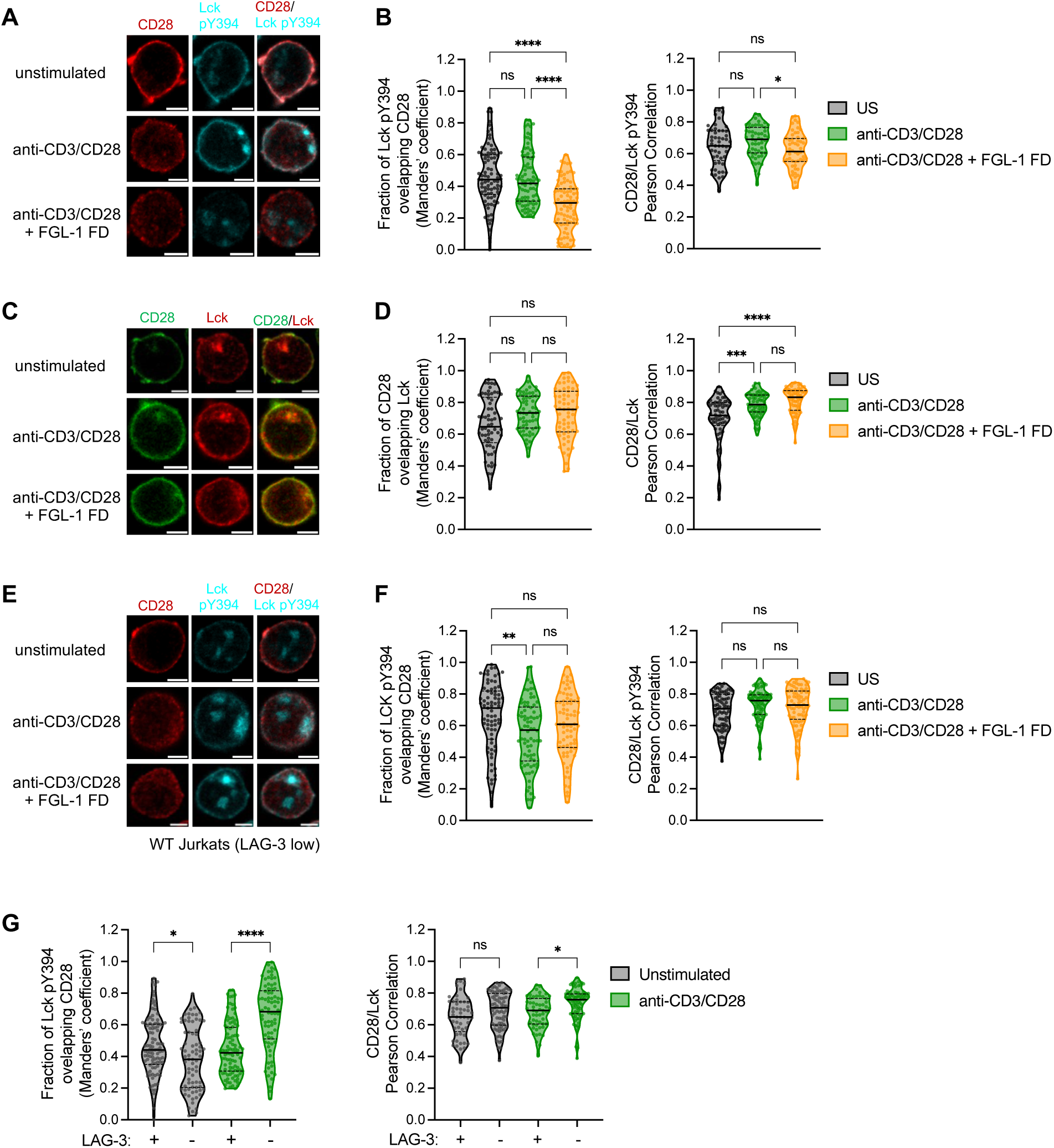
LAG-3/FGL-1 interaction downregulates CD28 and active Lck (pY394) colocalization in activated Jurkat T cells. (**A**) Representative fixed-cell confocal microscopy images showing colocalization of CD28 (red) and Lck (pY394, cyan) in unstimulated, anti-CD3/CD28-stimulated, and anti-CD3/CD28 +/− FGL-FD stimulated LAG-3+ Jurkat cells. Scale bar: 5 µm. **(B)** Violin plots quantifying CD28 and Lck pY394 colocalization using Manders’ overlap coefficient (MOC) and Pearson’s correlation coefficient (PCC) (n ≥ 50). **(C)** Representative confocal images showing colocalization of CD28 (green) and total Lck (red) in unstimulated, anti-CD3/CD28 +/− FGL-FD stimulated LAG-3+ Jurkat cells. Scale bar: 5 µm. **(D)** Colocalization analysis of CD28 and total Lck, measured by MOC and PCC (n ≥ 60). **(E)** Representative confocal images of CD28 (red) and Lck pY394 (cyan) in unstimulated, anti-CD3/CD28 +/− FGL-FD stimulated Jurkat cells. Scale bar: 5 µm. **(F)** Colocalization analysis of CD28 and Lck pY394, determined by MOC and PCC (n ≥ 75). **(G)** Comparison of MOC and PCC values for CD28 and Lck pY394 colocalization between unstimulated and stimulated LAG-3+ Jurkat cells compared to WT (LAG-3 low) Jurkat cells (n ≥ 75). Median values are shown in solid black lines; interquartile ranges as dashed black lines. Statistical analysis was performed using Kruskal-Wallis ANOVA test, followed by Dunnett’s post hoc tests. **** p < 0.0001, *** p < 0.001, ** p < 0.01, * p < 0.05, ns = not significant (p > 0.05).

In contrast, colocalization analyses revealed no differences in CD28 and total Lck overlap in Jurkat cells stimulated with or without FGL-1 FD (**Fig 6 C, D**), indicating that LAG-3/FGL-1-mediated inhibition specifically targets the active form of Lck. To confirm the specificity of this effect to the LAG-3/FGL-1 interaction, we examined the WT Jurkat cells (LAG-3 low). In these LAG-3-cells, T cell activation significantly increased CD28 and Lck pY394 colocalization, and importantly, FGL-1 FD treatment did not alter this interaction (**Fig 6 E, F**). Moreover, the proportion of Lck pY394 colocalizing with CD28 was significantly lower in LAG-3+ cells (MOC median: 42%) compared to LAG-3^−^ cells (MOC median: 68%) (**Fig 6 G**). This finding is consistent with previous studies describing a ligand-independent inhibitory effect of LAG-3.^31^ Notably, we observed no differences in CD28/LAG-3 colocalization in Jurkat LAG-3+ cells stimulated in the presence or absence of FGL-1 (**Fig. S10**), indicating that the enhanced proximity of LAG-3 and CD28 is independent of LAG-3 ligand binding. This suggests that FGL-1 binding specifically triggers the LAG-3 inhibitory mechanism and augments its effect beyond increased proximity due to the constitutive TCR/CD3–LAG-3 association.

To further substantiate these findings, we conducted additional colocalization analyses in primary T cells pre-activated to induce LAG-3 expression. Across three healthy donors we observed a broad range of MOC values for CD28 and Lck pY394 colocalization in LAG-3+ cells (**Fig 7 A, B**). This variability likely reflects differences in CD28 and LAG-3 expression among cells, as also evidenced by flow cytometry data (**Fig. S11**). Despite this heterogeneity, stimulation in the presence of FGL-1 FD consistently resulted in reduced CD28 and Lck pY394 colocalization compared to stimulation without FGL-1 FD. Collectively, these data suggest that LAG-3/FGL-1 interaction disrupts Lck pY394 binding and phosphorylation of CD28, thereby suppressing CD28-mediated costimulatory signaling.

**Figure 7.**
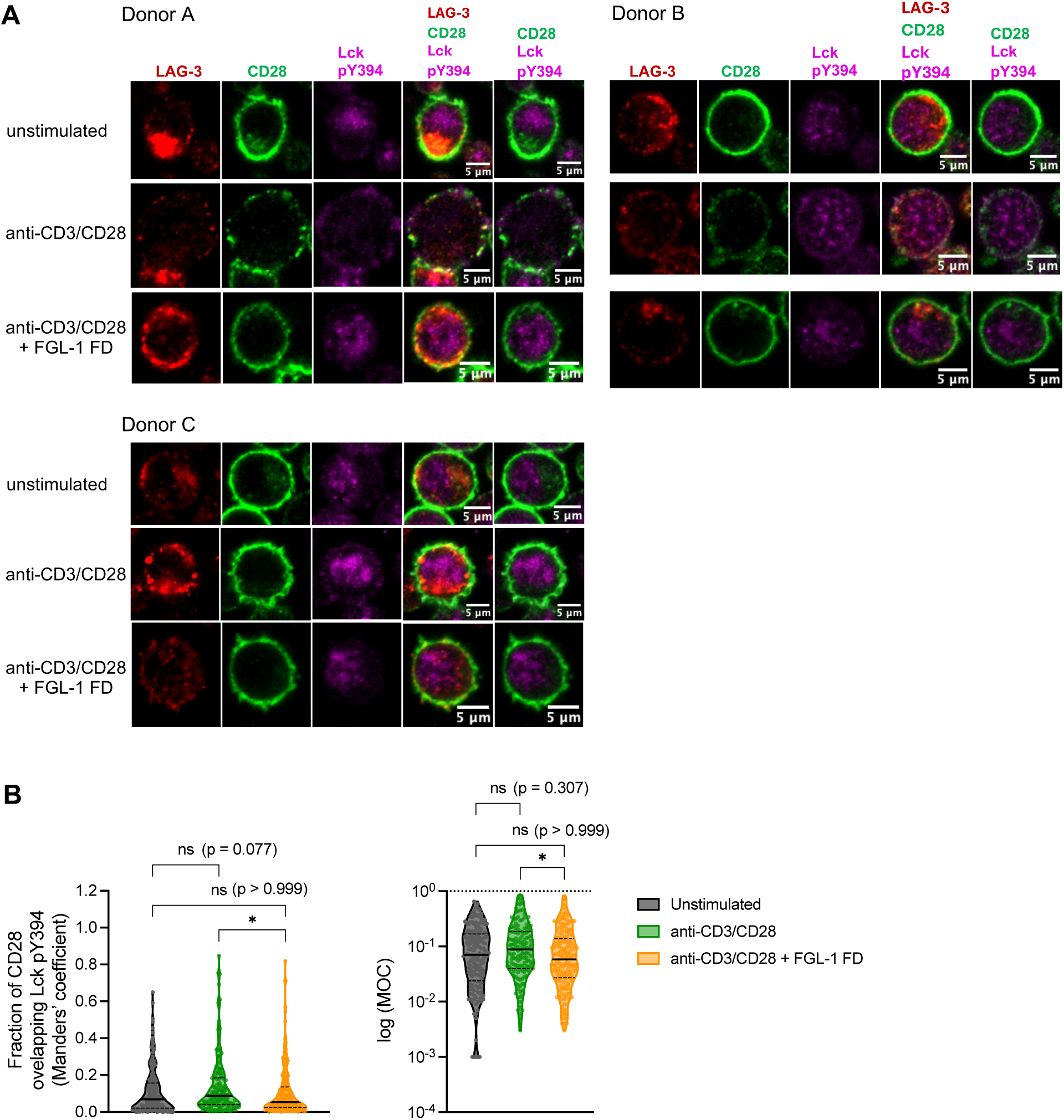
Reduced colocalization of CD28 and Lck pY394 in activated primary T cells. (**A**) Representative confocal images showing colocalization of CD28 and active Lck (pY394) in primary T cells from three healthy donors, pre-activated to induce LAG-3 expression. Images display LAG-3+ CD3+ cells from three healthy donors under unstimulated, stimulated with anti-CD3/CD28 +/− FGL-FD. Scale bar: 5 µm. **(B)** Quantitative colocalization analysis of CD28 and Lck pY394 expressed as Manders’ Overlap Coefficient (MOC) values and log (MOC), in CD3+/LAG-3+ cells across three donors (3 donors x n ≥ 50 per condition). Median values are shown as solid black lines, and interquartile ranges as dashed black lines. Statistical significance was assessed using Kruskal-Wallis ANOVA, followed by Dunnett’s post hoc tests. **** p < 0.0001, *** p < 0.001, ** p < 0.01, * p < 0.05, ns = non-significant (p > 0.05).

## DISCUSSION

Despite extensive research, the precise mechanisms by which LAG-3 regulates TCR/CD3 and CD28 signaling remain incompletely understood. Our study clarifies the inhibitory role of the LAG-3/FGL-1 interaction in modulating T cell activation through these pathways. We demonstrated that recombinant FGL-1 FD binds specifically to LAG-3 and markedly suppresses NF-κB induction, IL-2 secretion, and the expression of key T cell activation markers, including PD-1, CD69, CD25, ICOS, 4-1BB, and OX40. The specificity of the LAG-3/FGL-1 interaction was rigorously confirmed through binding and functional assays, which showed minimal impact on T cell activation in WT Jurkat cells expressing low levels of LAG-3, underscoring the targeted nature of LAG-3/FGL-1-mediated inhibition.

Additionally, our findings build on prior work by demonstrating that FGL-1 oligomerization is a key driver of high-affinity LAG-3 engagement and consequent LAG-3-mediated T cell suppression. ^17,18,30^ Structural and biophysical analyses demonstrate that dimerization of the FGL-1 FD, stabilized by an interchain disulfide bond, enhances LAG-3 binding affinity compared to the monomeric form. Multimerized FGL-1, generated via biotinylation and streptavidin conjugation, mimics this avidity effect, promoting stable receptor interaction. Functional assays in LAG-3+ Jurkat cells confirm the specificity and potency of this mechanism, supporting a model in which FGL-1 oligomerization facilitates efficient LAG-3 engagement. Thus, these results clarify the molecular basis of FGL-1/LAG-3 binding and highlight FGL-1 oligomerization as a potential therapeutic target.

Nonetheless, our findings contrast with recent studies that question FGL-1 as a functional ligand of LAG-3, instead proposing that pMHC II^18^ or ligand-independent LAG-3 suppresses TCR signaling.^31^ These reports do not exclude an inhibitory role of LAG-3/FGL-1 interaction on TCR signaling; rather, they suggest that previous studies may not have evaluated this effect under optimal conditions. In our work, we directly addressed this effect by co-immobilizing FGL-1 FD and TCR/CD3-stimulating antibodies on the same beads, ensuring that the LAG-3/FGL-1 interaction occurred concomitantly with T cell activation and enabling a direct assessment of its impact on T cell activation. Supporting this complexity, Jiang et al. recently showed that ligand engagement triggers LAG-3 ubiquitination, releasing its cytoplasmic tail from the membrane and activating the LAG-3 inhibitory pathway.^30^ They reported reduced LAG-3 ubiquitination upon FGL-1-Fc (both soluble and membrane-bound) binding compared to pMHC II binding; however, both ligands resulted in similarly suppressed IL-2 production. These findings suggest, despite distinct molecular effects, both FGL-1 and pMHC II ultimately converge on activating the LAG-3 inhibitory pathway.

Our study demonstrates that LAG-3/FGL-1 engagement broadly suppresses T cell activation by dampening phosphotyrosine signaling across both TCR and CD28 pathways. Using advanced SH2 superbinder–enrichment phosphoproteomics, we revealed that the LAG-3/FGL-1 interaction leads to reduced phosphorylation at key sites on CD3, ZAP-70 and CD28, indicating a coordinated regulation of two essential axes for T cell activation. These findings, corroborated by Western blot and flow cytometry in both Jurkat and primary T cells, extend previous findings by elucidating the integrated regulation of TCR and costimulatory signaling.

Moreover, our data indicate that LAG-3/FGL-1 interaction suppresses CD28 costimulatory signaling on a similar timescale as TCR/CD3 signaling, occurring within 1 minute. This observation aligns with previous reports demonstrating spatiotemporal colocalization and signaling synergy, between the TCR/CD3 complex and CD28, which together enhances T cell signal transduction.^48,49^ Previous studies have shown that LAG-3 clusters and associates with the TCR/CD3 complex, exerting its inhibitory function by displacing Lck from CD4 and CD8 co-receptors in a ligand-independent manner.^31,46^ Our confocal imaging analysis revealed a significant increase in LAG-3 and CD28 colocalization following T cell activation, suggesting enhanced spatial proximity and potential interaction. However, in the presence of FGL-1 FD, we observed a marked reduction in CD28 and active Lck colocalization, indicating that LAG-3/FGL-1 engagement disrupts the recruitment of CD28 by Lck, uncoupling critical positive feedback necessary to sustain T cell activation. This inhibitory mechanism appears to be highly specific: WT Jurkat cells expressing little to no LAG-3 did not exhibit decreased CD28–Lck pY394 colocalization upon FGL-1 FD stimulation. Our findings demonstrate that this inhibition occurs independently of CD4 coreceptor-bound Lck, indicating that LAG-3 targets both free and membrane-associated Lck. This insight expands current mechanistic understanding of LAG-3 signaling beyond models that predominantly focus on coreceptor-associated Lck, suggesting a more comprehensive spatial and biochemical modulation of T cell signaling by LAG-3. Importantly, the reduction in CD28-active Lck association in LAG-3+ cells was consistently observed across various experimental conditions and cell types, including primary T cells exhibiting heterogenous CD28 and LAG-3 expression levels. The reproducibility of this effect underscores the robustness and physiological relevance of our findings. These results parallel a recently proposed LAG-3 inhibitory mechanism whereby LAG-3 downregulates CD3ε/Lck association,^32^ suggesting that LAG-3 primarily targets Lck binding and recruitment within the immunological synapse.

Our study extends the current understanding of LAG-3 function by demonstrating that the LAG-3/FGL-1 interaction directly regulates CD28 costimulatory signaling, offering a new perspective on LAG-3-mediated immunosuppression. Notably, suppression of CD28 phosphorylation by LAG-3 parallels one of the proposed the inhibitory mechanism of PD-1, which attenuates T cell activation by recruiting the SH2-containing tyrosine phosphatase-2 (SHP-2) to primarily dephosphorylate CD28,^50^ as well as CD3ζ and ZAP-70.^51^ This mechanistic similarity underscores the critical role of CD28 in T cell activation and highlights a shared axis of regulation among key coinhibitory receptors.

Furthermore, CD28 expression has emerged as a key determinant of responsiveness to PD-1 blockade. Clinical studies have shown that CD28 signaling is essential for the reinvigoration and proliferation of exhausted of CD8+ T cells following PD-1 blockade.^52–55^ Our findings suggest that LAG-3 may exert a comparable inhibitory effect by suppressing CD28 signaling, potentially contributing to resistance observed in patients receiving PD-1 monotherapy. This supports the rational for dual blockade of PD-1 and LAG-3, a strategy currently under investigation, aimed at enhancing antitumor immunity and improving outcomes in cancer patients.^56–58^ Additionally, our results also suggest that therapeutic strategies enhancing CD28-Lck interactions or preventing their disruption may synergize with PD-1/LAG-3 blockade to improve antitumor immunity. Further investigation into spatial regulation of kinase-receptor interactions at the immunological synapse could uncover new avenues to overcome resistance mechanisms limiting immunotherapy efficacy.

While our proposed LAG-3/FGL-1 inhibitory mechanism is supported by multiple biochemical and cellular imaging approaches, certain aspects of the study, particularly those involving primary cells, were subject to variability. This likely stems from heterogeneous expression levels of CD28 and LAG-3 across individual cells, influencing the consistency of colocalization and phosphorylation measurements. Future studies should consider standardizing activation conditions or stratifying cells based on expression profiles to improve reproducibility. Additionally, our use of fixed-cell confocal microscopy, while valuable for spatial resolution, does not capture dynamic molecular interactions. Live-cell imaging approaches in physiologically relevant immune contexts will be instrumental in elucidating the real-time impact of LAG-3/FGL-1 engagement on T cell signaling. Moreover, the molecular determinants governing LAG-3 targeting of free versus coreceptor-bound Lck pools warrant deeper biochemical and structural characterization.

Despite these limitations, the convergence of data across complementary analyses strongly supports a model in which the LAG-3/FGL-1 interaction suppresses CD28 costimulatory signaling, through the disruption of CD28-Lck recruitment and phosphorylation. These results point to a broader inhibitory role for LAG-3, targeting multiple nodes within the TCR/CD3 signaling cascade to regulate T cell activation. These insights not only deepen our understanding of T cell inhibition but also identify the LAG-3/FGL-1 axis as a promising target for enhancing immune-based therapies. Further investigation into this pathway in disease models will be critical for translating these findings into clinical applications. Such efforts have the potential to advance not only cancer immunotherapy but also therapeutic strategies for autoimmune and infectious diseases.

### Resource Availability

Further information and requests for resources and reagents should be addressed to the lead contact, Michelle Krogsgaard.

### Data Availability

X-ray crystallography data for the structures of ligand-free mutant Src SH2 SB and mutant Src SH2 SB bound to O-Phospho-L-tyrosine have been deposited to the Protein Data Bank under the accession codes 8VCF and 8VCG, respectively.

## Acknowledgements

We thank Y. Velmurugu, A. Natarajan, and K. Ruggles (NYU Grossman School of Medicine) for helpful discussions and feedback on our manuscript; J. Wang and J. Du (NYU Grossman School of Medicine) for generously providing the LAG-3+ NF-κB::eGFP reporter Jurkat E6.1 T cells; and Y. Deng and M. Cammer (NYU Grossman School of Medicine) for overseeing the use of the Microscopy Laboratory. Part of this research was performed at the Advanced Photon Source, a U.S. Department of Energy (DOE) Office of Science user facility operated for the DOE Office of Science by Argonne National Laboratory under Contract No. DE-AC02-06CH11357. Mass spectrometry experiments were supported in part by NYU Langone Health and the Laura and Isaac Perlmutter Cancer Center support grant P30CA016087 from the National Cancer Institute. The NYU Microscopy Center is also partially supported by the Cancer Center Support Grant P30CA016087 at the Laura and Isaac Perlmutter Cancer Center. This work was supported by NIH grant NCI R01 CA243486 (M.K.) and Merck Oncology Translational Science Program (OTSP) grants 57570 and 58166 (M.K.).

## Author Contributions

M.K. led project administration and supervision. Experimental conceptualization and design were developed by S.N., Y.P., and M.K. Methodology development, data curation, and formal analysis were performed by S.N., E.K., Y.P., and B.U. Experimental investigation was conducted by S.N., A.H., E.K., N.O., and Y.P. Validation and visualization were carried out by S.N. Funding acquisition was managed by B.U. and M.K. Manuscript was written by S.N., Y.P., and M.K.

## MATERIALS AND METHODS

### Cell lines and PBMC sample processing

LAG-3+ NF-κB::eGFP reporter Jurkat E6.1 T cells were a gift from Jun Wang (NYU). WT Jurkat (Clone E6-1) cell line was obtained from the American Type Culture Collection (ATCC). Jurkat cells and primary human T cells were cultured in RPMI 1640 medium supplemented with 10% fetal bovine serum (FBS), 2 mM L-glutamine, 1X penicillin/streptomycin), maintained at in a 37°C in humidified atmosphere containing 5% CO2.

Peripheral blood leukopaks from healthy donors were obtained from New York Blood Center Enterprises. Peripheral blood mononuclear cells (PBMCs) were isolated by Ficoll-Hypaque (GE Healthcare) density gradient centrifugation. The PBMC layer was carefully collected, pelleted, and the supernatant was aspirated without disturbing the cell pellet. PBMCs were then resuspended in human serum (HyClone) containing 10% dimethyl sulfoxide (DMSO) and aliquoted into cryovials with freezing medium composed of FBS and 10% DMSO (Sigma). Cryovials were placed in CoolCell freezing containers at −80 °C for 24 hours prior to transfer to liquid nitrogen (−140 °C). For culture, PBMCs were rapidly thawed, resuspended in culture medium, and maintained at 37°C with 5% CO_2_.

### Protein sequences, expression, refolding and purification

Amino acid sequences for human FGL-1 Fibrinogen Domain (FD) (residues 74-306), and Src SH2 domain (residues 151-248) were obtained from the UniProt.^59^ Corresponding codon-optimized cDNA for bacterial expression were synthesized and cloned into pET 30a+ (FGL-1) and pET 28a (Src) and vectors (Genewiz). The FGL-1 FD construct included a C-terminal Avi-tag (GLNDIFEAQKIEWHE) for site-specific biotinylation.^60^ The Src SH2 protein construct incorporated mutations (Thr183Val, Ser188Ala, Lys206Leu, Cys241Ser, Cys248Ser) to enhance phosphotyrosine binding affinity. Site-directed mutagenesis was performed using the Q5® Site-Directed Mutagenesis Kit (New England Biolabs) following the manufacturer protocols; all constructs were verified by DNA sequencing.

FGL-1 FD plasmids were transformed into *E. coli* BL21 (DE3), which were cultured in Terrific Broth supplemented with 50 ug/mL kanamycin at 37 °C OD600 reached 0.6. Protein expression was induced with IPTG (Thermo Scientific) and incubated at 18 °C for 16 hours. FGL-1 FD was expressed as insoluble inclusion bodies (IBs). Cells were lysed by lysozyme treatment, osmotic shock, and freeze-thaw cycles. IBs were isolated by centrifugation and washed multiple times with 0.5–2.0% Triton X-100 to remove contaminants. Purified IBs were solubilized in 6 M guanidine hydrochloride, insoluble debris removed by centrifugation, and protein refolded by rapid dilution into refolding buffer containing 0.5 M L-arginine, 0.1 M Tris-HCl (pH 8), 2 mM EDTA, 5 mM reduced glutathione, and 0.5 mM oxidized glutathione. Refolded protein was dialyzed against 10 mM Tris-HCl (pH 8) to remove denaturants. Final purification was performed by size exclusion chromatography (SEC) on Superdex 200 and Superdex 75 columns (Cytiva Life Sciences) in 10 mM Tris-HCl (pH 8) with 0.1 M NaCl using an ÄKTA Pure25 system. For monomeric FGL-1 FD, refolding and purification were conducted in EDTA-free buffer supplemented with 2 mM CaCl_2_ to stabilize the protein.

Mutant Src SH2 SB domain were expressed as soluble hexahistidine-tagged proteins. Transformed colonies were cultured in Luria Broth supplemented with 50 ug/mL Kanamycin at 37 °C. Protein expression was induced with 1 mM IPTG for 4 hours, and cells were harvested by centrifugation and resuspended in buffer containing 25 mM HEPES (pH 7.0), 0.5 M NaCl, and 25% sucrose, subjected to two freeze-thaw cycles, treated with lysozyme (0.2 mg/mL) at 37°C for 15 minutes, then incubated with DNAse (Thermo Fisher Scientific) and MgCl₂ to degrade nucleic acids. Clarified lysates were purified by Ni-NTA affinity chromatography (GE Healthcare), followed by SEC on a Superdex 75 column (Cytiva Life Sciences) equilibrated with 25 mM HEPES-NaOH (pH 7.0) and 0.1 M NaCl using an ÄKTA Pure25 system.

### Protein Biotinylation

Recombinant *BirA* enzyme was expressed in *E. coli* and purified in house as previously described.^61^ Avi-tagged FGL-1 FD proteins were biotinylated using Avidity *BirA* Biotin Ligase kit, following manufacturer instructions. Proteins (40 μM final) were incubated with 0.05 M bicine buffer (pH 8.3), 10 mM ATP, 10 mM MgOAc, and 50 μM D-biotin. BirA enzyme was added at a 2.5 ug of Bir A per 10 nmol Avi-Tag substrate. Reactions proceeded at 37°C for 2.5 – 3 hours. Biotinylated proteins were buffer exchanged into 10 mM Tris-HCl (pH 8), 0.1 M NaCl using Zeba Spin Desalting Columns (Thermo Fisher Scientific). Successful biotinylation was confirmed by SDS-PAGE gel-shift assay in the presence of excess streptavidin.

### Biolayer Interferometry

Binding between FGL-1 FD and LAG-3 were measured BLI Octet RED96 instrument (Pall ForteBio) using streptavidin (SA)-conjugated biosensors, as per manufacturer’s instructions and published protocols.^62,63^ All experiments were performed at 25°C in HST buffer (10 mM HEPES, pH 7.5; 100 mM NaCl; 0.01% Tween-20). Biotinylated FGL-1 FD (1 μg/mL) was immobilized on SA sensors to a loading response of ∼0.25 nm. After a 2-minute baseline equilibration, sensors were exposed to varying LAG-3 concentrations (0 – 480 nM) for 3 minutes association and a 4-minute dissociation step in buffer. Parallel experiments minimized batch effects. Data were analyzed using fitted using Octet® Data Analysis software version 9.1 to derive kinetic parameters and equilibrium dissociation constants (K_D_).

### Protein Crystallization, Structure Solution and Refinement

Proteins were crystallized by sitting-drop vapor diffusion methods with a 1:1 (v/v) protein-to-precipitant ratio. Initial crystallization screens utilized MCSG kits (Anatrace) in 96-well Intelli-plates (Art Robbins Instruments), set up by a Mosquito robot (SPT Labtech) and incubated at 18°C. Crystals appeared within 3–10 days and were manually optimized.

FGL-1 FD dimer (without Avi-tag) crystallized in 0.1 M HEPES (pH 7.5), 10% PEG 8000 (w/v), and 8% ethylene glycol (v/v) at 10 mg/ml protein concentration at 18°C. Plate-like crystals formed within 2–5 days and were flash-frozen in liquid nitrogen after brief soaking in precipitant supplemented with 30% ethylene glycol.

Src SH2 domain crystals grew in 0.2 M ammonium fluoride and 20% PEG 3350 over 7 days. Optimization involved mixing 0.5 μL of 15 mg/mL protein with varying precipitant volumes (0.5– 1.5 μL) in 24-well plates. pY-bound Src SH2 crystals were obtained by soaking apo crystals in 10 mM O-phospho-L-tyrosine. Crystals were cryoprotected and flash-frozen in liquid nitrogen prior to X-ray diffraction.

Diffraction data were collected at 100°K at the APS beamline 19-ID (FGL-1 FD) and 19-BM (Src SH2 SB). Data integration and processing was performed HKL2000 (19-ID) or HKL3000 (19-BM) and scaled using AIMLESS (CCP4 suite).^64–66^ Structures were solved by molecular replacement (MR) using PHASER^67^ with PDB 1FZD (FGL-1 FD) or 4F5B (Src SH2) as a search models. Refinement was done with REFMAC^68^ and manual rebuilding in COOT.^69^ Models were validated using Sfcheck, Procheck software in CCP4, Molprobity, and PDB validation tools. Data collection and refinement statistics, along with PDB accession codes, are provided in **Supplementary Tables 1 and 2**. Structural figures were prepared with PyMOL (Schrödinger, L. & DeLano, W., 2020). *PyMOL*, Available at: http://www.pymol.org/pymol).

### Assembly of FGL-1 FD Streptavidin-Phycoerythrin tetramers

Biotinylated FGL-1 FD was tetramerized with Streptavidin-Phycoerythrin (SAPE; Thermo Scientific) for flow cytometry. Streptavidin-PE was added dropwise at 4 x molar excess over biotinylated protein, in aliquots of 1/10 volume every 3 minutes at room temperature in the dark. After the final addition, the mixture was incubated for 10 min at room temperature. Tetramers were buffer-exchanged into 10 mM Tris-HCl (pH 8), supplemented with 2 mM CaCl_2_ for FGL-1 FD monomers, using Amicon Ultra centrifugal filters (Millipore). Tetramers were titrated onto LAG-3– expressing Jurkat cells, and binding was assessed by flow cytometry.

### T cell activation using M280 Streptavidin Dynabeads

M280 streptavidin Dynabeads (Thermo Scientific) were coated with biotinylated anti-human CD3 antibody (clone OKT3, Invitrogen), anti-human CD28 antibody (clone CD28.2, Invitrogen), and recombinant FGL-1 FD at a molar ratio of 1:1:5 (anti-CD3:anti-CD28:FGL-1 FD). Controls lacking FGL-1 FD included biotinylated mouse IgG isotype antibody to normalize biotin content. Jurkat T cells or primary human T cells were stimulated with coated beads at a 1:5 cell-to-bead ratio. For short-term stimulation, activation was halted by the addition of cold 1× PBS (3× culture volume). For overnight stimulation, culture supernatants were harvested, and IL-2 secretion was quantified using the Human IL-2 ELISA MAX™ kit (BioLegend) per manufacturer’s instructions. Jurkat T cell activation was additionally assessed by measuring NF-κB::eGFP expression using flow cytometry.

### Flow cytometry

Jurkat T cells or primary PBMCs were washed twice with 1× PBS to remove residual media and serum. For surface staining, cells were incubated with a viability dye for 15 minutes in the dark, washed, and then stained with fluorophore-conjugated antibodies for 30 minutes. For phospho-flow assays, cells were fixed immediately after stimulation and permeabilized using the BD Pharmingen™ Transcription Factor Phospho Buffer Set (BD Biosciences) following manufacturer’s instructions. Simultaneous intracellular and surface staining was performed concurrently in the dark for 1 hour. Following staining, cells were washed and resuspended in FACS buffer (1× PBS with 1% FBS) prior to acquisition on a Cytek Aurora flow cytometer (Cytek). The following antibodies were used:

T cell activation markers:

BUV563-conjugated anti-CD25, BV421-conjugated anti-CD279 (PD-1), PerCP-eFluor 710-conjugated anti-CD278 (ICOS), BV650-conjugated anti-CD69, BUV805-conjugated anti-CD134 (OX40), PECy5-conjugated anti-CD137 (4-1BB)

Phospho-specific antibodies:

SparkBlue 550-conjugated anti-CD3, SparkViolet 538-conjugated anti-CD4, Brilliant Violet™ 570-conjugated anti-CD8, Brilliant Violet™ 421-conjugated anti-CD223(LAG-3), Brilliant Blue 700-conjugated anti-CD279 (PD-1), Brilliant Violet™ 786-conjugated anti-CD28, PE-Cy7-conjugated anti-anti-ZAP70 (pY319), PE-CF594-conjugated anti-Akt (pS473), Alexa Fluor 700-conjugated anti-ERK 1/2 (pT202/pY204), Alexa Fluor647-conjugated anti-pPDK1 S241, PE-conjugated anti-Lck (pY394).

Data analysis was performed using FlowJo V10 Software.

### Western Blotting

Cell pellets were lysed in ice-cold buffer (20 mM Tris-HCl pH 7.5, 150 mM NaCl, 1% n-dodecyl-β-d-maltoside, 1 mM Na_3_VO_4_, 1x complete protease inhibitor cocktail (Roche) and 2% phosphatase inhibitor mixture 2 (Sigma-Aldrich) for 30min at 4°C. Lysates were clarified by centrifugation, separated by SDS-PAGE and transferred to nitrocellulose membranes. Membranes were blocked in Intercept^®^ (TBS) Blocking Buffer (Licor), then probed overnight with the following antibodies Lck Y394 and total (Cell signaling, Santa Cruz), Zap70 Y319 and total (Cell signaling, Santa Cruz), CD28 Y191(Cell signaling), Erk1/2 T202/Y204 and total (Cell signaling), CD3ζ Y142 (Cell signaling). Detection was performed using IRDye-conjugated secondary fluorescent anti-Rabbit and anti-Mouse antibodies and visualized on the Odyssey CLx Imaging system.

### Immunofluorescence Imaging and Colocalization Analysis

Jurkat T cells or primary T cells were added to poly-L-lysine-coated chambered coverslips (Lab-Tek, Thermo Fisher) pre-coated with 5 μg/mL human anti-CD3 antibody (clone SK7, Miltenyi Biotec), 5 μg/mL anti-CD28 antibody (clone L293, Miltenyi Biotec), and 20 μg/mL FGL-1 FD or human IgG isotype control antibody in 1× PBS. After 5 minutes at 37°C, cells were fixed with 2 % paraformaldehyde for 10 mins and permeabilized in 0.05% digitonin. Blocking was performed with 2% goat serum for 1 hour. Cells were stained with primary antibodies in 10% FBS/PBS) for 1 hour (or overnight at 4°C), washed, and then incubated with fluorophore-conjugated secondary antibodies. DAPI was used for nuclear staining.

Antibody-fluorophore combinations for imaging included:

**Figure 5**: Alexa Fluor 568 and 647 conjugates (Invitrogen) were used to detect LAG-3 and CD3 or CD28, respectively. **Figure 6**: Alexa Fluor 568 and 647 (Invitrogen) were used to detect CD28 and pLck-Y394, respectively. **Figure 7**: Alexa Fluor 488, 568 and 647 (Invitrogen) were used to detect CD28, LAG-3 and pLck-Y394, respectively.

Images were aquired using a Zeiss LSM 880 confocal microscope with a 63X/1.40 oil objective using. Colocalization analysis was performed in Fiji software.^70^ Colocalization per cell was quantified by Manders’ overlap coefficient (MOC) and Pearson’s correlation coefficient (PCC) using the Just Another Colocalization (JACoP) plugin.^71,72^

### SH2 superbinder resin preparation

Src SH2 superbinder (SB) was covalently conjugated to CNBr-activated Sepharose resin (Sigma). The resin was activated and washed with cold 1 mM HCl, then equilibrated in coupling buffer (100 mM NaHCO₃, pH 8.3, 500 mM NaCl). SB was buffer exchanged and added to beads at a 1:100 ratio (w/w) overnight at 4°C. Unbound sites were blocked (0.1 M Tris-HCl, 0.75 M glycine, pH 8.0). Beads were washed and stored in immunoaffinity purification (IAP) buffer (50 mM Tris-HCl, pH 7.2; 50 mM NaCl; 10 mM Na₂HPO₄) + 0.02% sodium azide at 4°C.

### Proteomics sample preparation and pY peptide enrichment

Jurkat LAG-3+ cells were left unstimulated or stimulated with anti-CD3/CD28 Abs +/− FGL-1 FD, then lysed in urea buffer (8 M urea, 50 mM Tris-HCl, pH 8, plus protease and phosphatase inhibitors). Proteins were reduced (5 mM TCEP) and alkylated (10 mM CAA) at 56°C for 1 hour. clarified, and protein concentrations were quantified by absorbance at 280nm using a Little Lunatic device (Unchained Labs). Samples were diluted 6-fold in 20 mM Tris (pH 8) and digested overnight with trypsin (1:100 enzyme-to-protein ratio). Peptides were acidified (1% FA), desalted, and lyophilized. Peptides were reconstituted in cold IAP buffer and incubated overnight with the SH2 SB beads at 4 °C. Enriched peptides were eluted (0.1 % TFA, 30% ACN), lyophilized, and resuspended for LC-MS/MS.

### LC-MS/MS and DIA Acquisition

Peptides were separated on a Dr Maisch C18 AQ (150µm ID, 15cm) column using an EvosepOne LC^73^ system and analyzed on an Orbitrap HF-X mass spectrometer using 88 min extended Evosep method (SPD15) at a flowrate of 220 nl/min. Mass spectrometer was operated in data-independent acquisition mode (DIA)^74^ doing MS^2^ fragmentation across 22 m/z windows after every MS^1^ scan event (see table below).

| window | start_mz | end_mz | center | width |
| --- | --- | --- | --- | --- |
| 1 | 350 | 384 | 367 | 34 |
| 2 | 383 | 411 | 397 | 28 |
| 3 | 410 | 436 | 423 | 26 |
| 4 | 435 | 460 | 447.5 | 25 |
| 5 | 459 | 482 | 470.5 | 23 |
| 6 | 481 | 504 | 492.5 | 23 |
| 7 | 503 | 527 | 515 | 24 |
| 8 | 526 | 548 | 537 | 22 |
| 9 | 547 | 571 | 559 | 24 |
| 10 | 570 | 593 | 581.5 | 23 |
| 11 | 592 | 616 | 604 | 24 |
| 12 | 615 | 640 | 627.5 | 25 |
| 13 | 639 | 666 | 652.5 | 27 |
| 14 | 665 | 693 | 679 | 28 |
| 15 | 692 | 723 | 707.5 | 31 |
| 16 | 722 | 755 | 738.5 | 33 |
| 17 | 754 | 791 | 772.5 | 37 |
| 18 | 790 | 834 | 812 | 44 |
| 19 | 833 | 885 | 859 | 52 |
| 20 | 884 | 956 | 920 | 72 |
| 21 | 955 | 1071 | 1013 | 116 |
| 22 | 1070 | 1650 | 1360 | 580 |

High-resolution full MS spectra were acquired with a resolution of 120,000, an AGC target of 3e6, a maximum ion injection time of 60 ms, and a scan range of 350 to 1650 m/z. Following each full MS scan, 22 data-independent HCD MS/MS scans were acquired at a resolution of 30,000, an AGC target of 3e6, and stepped nices of 22.5, 25, and 27.5.

### Proteomics Data Processing

Data were processed in Spectronaut (https://biognosys.com/shop/spectronaut) using the directDIA workflow. Searches were performed against the human Uniprot database (http://www.uniprot.org/) with tryptic specificity (max 2 missed cleavages) concatenated with a list of common lab contaminants. Database search was performed in the integrated search engine Pulsar. Oxidation of methionine and phosphorylation on serine, threonine, and tyrosine residues were set as variable modifications; carbamidomethylation of cysteines was set as a fixed modification. The false discovery rate (FDR) was controlled at 1% at the peptide, protein, and site level. Phosphosite localization was set to >75% confidence. Protein quantification was performed on MS^2^ level using 3 most intense fragment ions per precursor. Label-free normalization across the runs were performed based on the median log2-transformed intensities of the peptide precursors in each run.

Subsequent data analysis steps were performed in either Perseus^75^ (http://www.perseus-framework.org/) or using R environment for statistical computing and graphics (http://www.r-project.org/).

### Statistical analyses

GraphPad Prism software was used for unpaired two-tailed Student’s t-tests, one-way ANOVA, and two-way ANOVA; and p values <0.05 were considered significant (*p<0.05, **p<0.01, ***p<0.001 and ****p<0.0001). Statistical enrichment of phosphorylation sites and pathways were determined on Perseus^76^ and Database for Annotation, Visualization and Integrated Discovery (DAVID) Bioinformatics Resources.^42,43^ For proteomics, statistical was assessed using Welch’s t-test or ANOVA, corrected by Benjamini-Hochberg False Discovery Rate (FDR) set to 5%. Error bars denote standard deviation (SD) unless otherwise indicated.

## Supplementary Figures

**Supplementary Figure 1.**
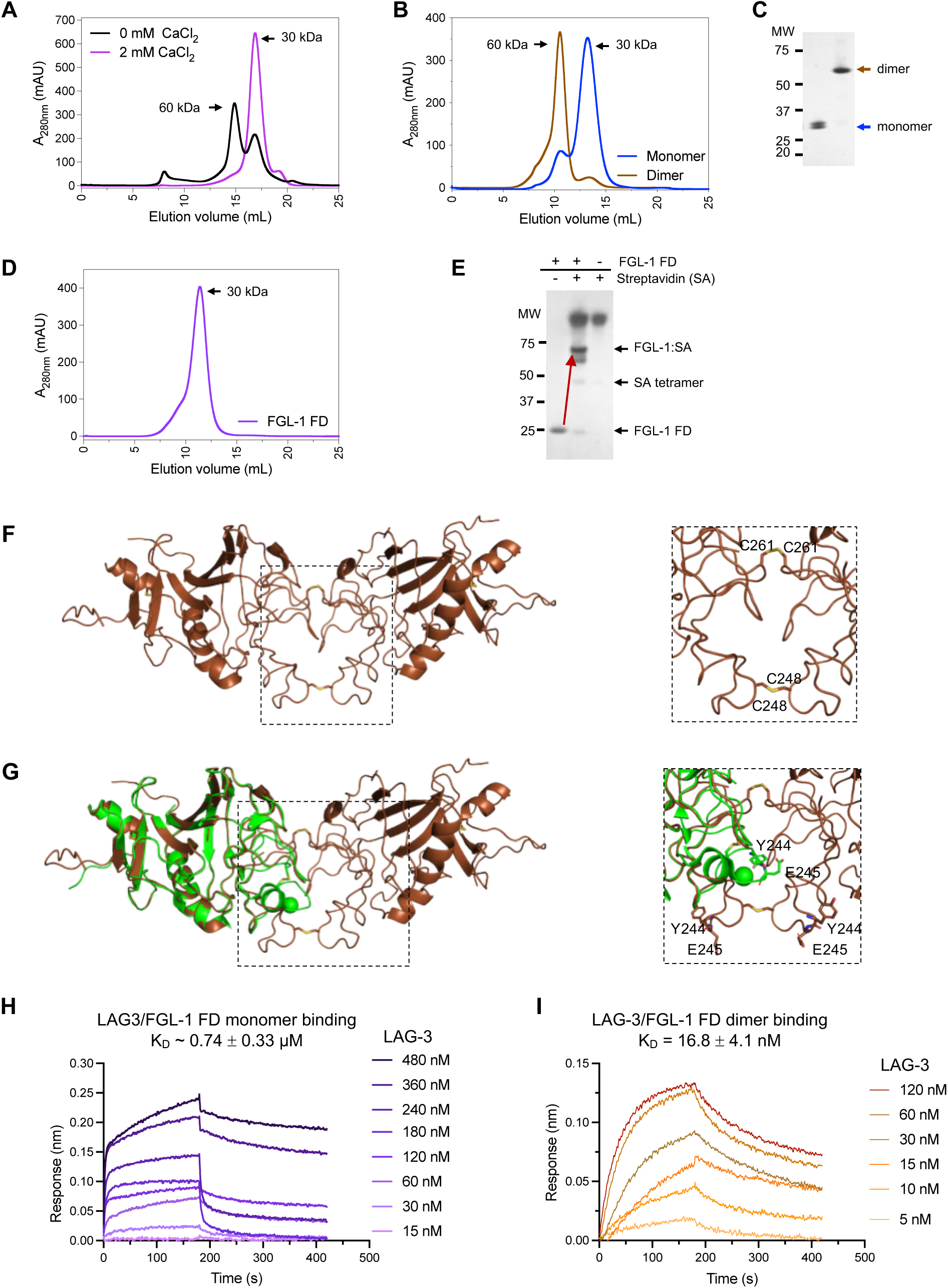
FGL-1 FD undergoes Ca^2+^-dependent conformational changes resulting in monomeric and dimeric states. (**A**) Superimposed Superdex 200 (S200) size exclusion chromatography (SEC) elution profiles of refolded FGL-1 FD +/− of 2 mM CaCl_2_. Arrows indicate elution volumes corresponding to monomeric and dimeric species. **(B)** FGL-1 FD monomeric and dimeric fractions from (**A**) were further purified on Superdex 75 (S75) Increase column. **(C)** SDS-PAGE of FGL-1 FD monomer and dimer fractions purified in (**B**). Arrows denote protein band at expected molecular weights. **(D)** S75 SEC elution profile of purified, refolded FGL-1 FD confirming separation of monomer and dimer states. **(E)** SDS-PAGE of biotinylated FGL-1 FD. The red arrow indicates a streptavidin-induced protein band shift, consistent with functional biotinylation. **(F)** High-resolution crystal structure of FGL-1 FD dimer, resolved at 1.74 Å, shows intermolecular disulfide bonds stabilizing the dimeric conformation. **(G)** Structural alignment of FGL-1 FD dimer (brown) and FGL-1 FD monomer (PDB: 7TZ2; magenta). The root mean square deviation (RMSD) = 0.397 Å. **(H, I** Bio-Layer Interferometry (BLI) sensograms of LAG-3 binding to FGL-1 FD monomer (**H**) and dimer (**I**). Dissociation constants (K_D_) were calculated using steady-state analysis

**Supplementary Figure 2.**
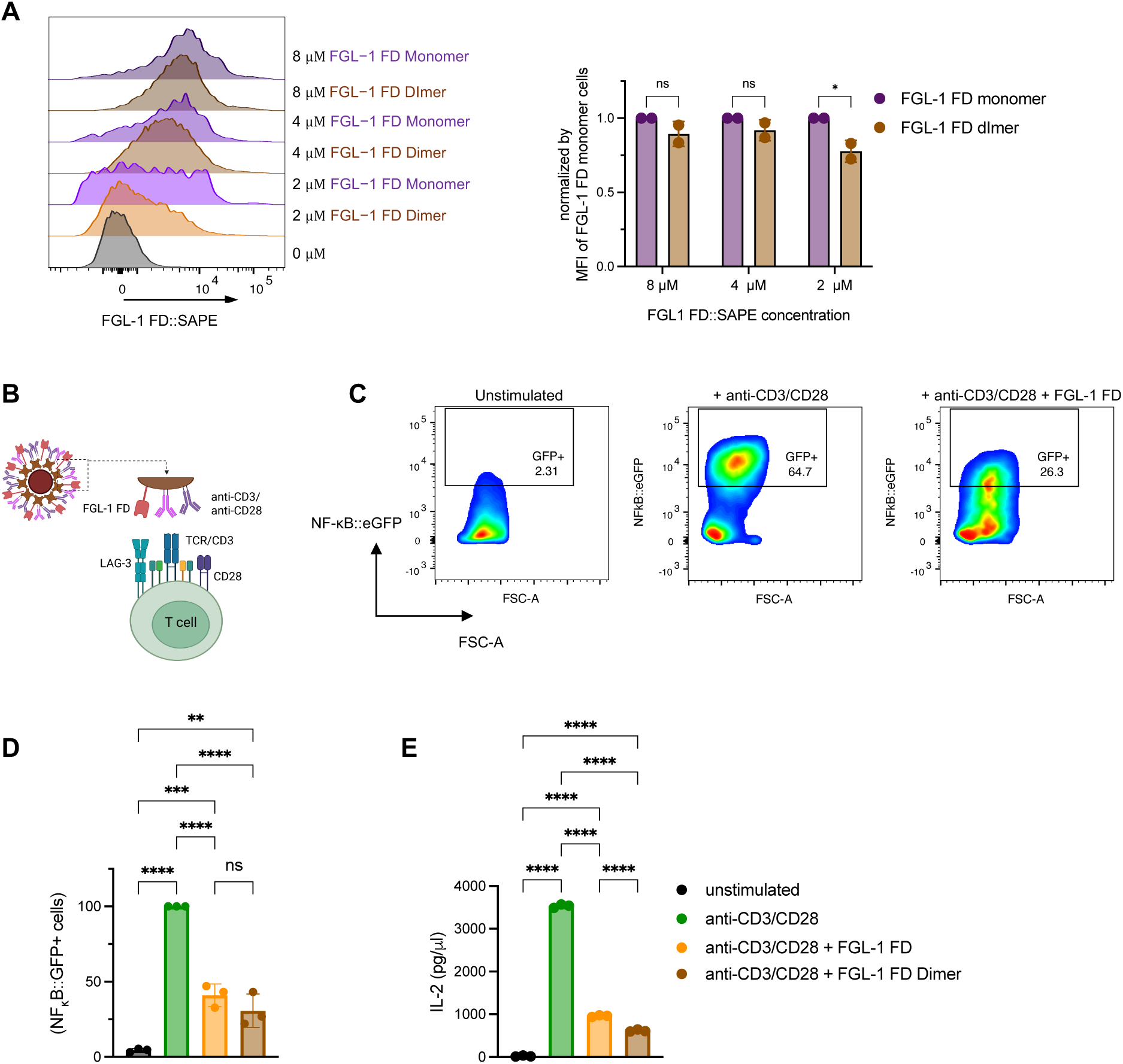
Functional validation of recombinant FGL-1 FD. (**A**) Flow cytometry histograms showing binding of FGL-1 FD monomer and dimer tetramer bindings to LAG-3+ Jurkat cells (n = 2). Data are presented as median fluorescent intensity (MFI), normalized by FGL-1 FD monomer tetramer binding at each concentration. **(B)** Schematic representation of LAG-3+ T cell activation using M280 streptavidin Dynabeads coated with anti-CD3 and anti-CD28 antibodies, and FGL-1 FD. **(C)** Representative flow cytometry plots showing NF-κB::eGFP expression in LAG-3+ Jurkat cells stimulated with anti-CD3/CD28 antibodies plus FGL-1 monomer, dimer, or mouse IgG1 isotype control (mIgG1)-coated beads. **(D)** Quantification of NF-κB::eGFP induction, shown as percentage activation normalized to MFI of anti-CD3/CD28 stimulated LAG-3+ Jurkat cells (100%) (n =3). **(E)** IL-2 production in unstimulated and stimulated LAG-3+ Jurkat cells under the same bead-based stimulation conditions. Statistical significance was determined using one-way ANOVA, Statistical significance was performed using one or two-way ANOVA, followed by Tukey or Šídák’s post hoc tests, respectively. Error bars represent standard deviation (SD). **** p < 0.0001, *** p < 0.001, ** p < 0.01, * p < 0.05, ns = not significant (p > 0.05).

**Supplementary Figure 3.**
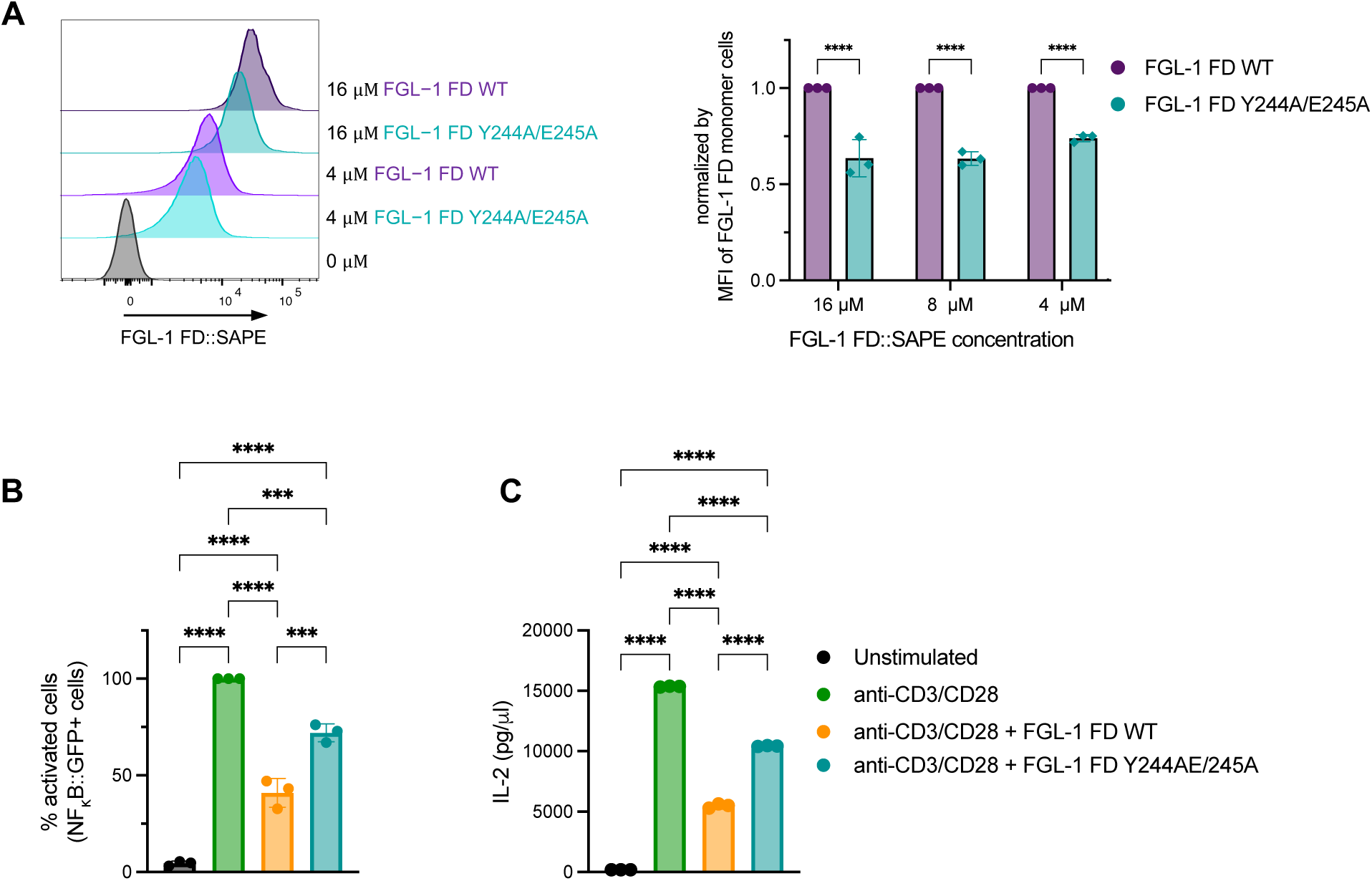
Functional validation of FGL-1 FD loss-of-function mutant, Y244A/E245A (putative LAG-3 interaction site). (**A**) Flow cytometry histograms showing binding of FGL-1 FD wild-type (monomer) and Y244A/E245A mutant tetramers to LAG-3+ Jurkat cells (n = 3). Data are presented as median fluorescent intensity (MFI) normalized to FGL-1 FD monomer tetramer binding at each concentration. **(B)** NF-κB::eGFP induction and **(C)** IL-2 production in LAG-3+ Jurkat cells stimulated for 24 hours with anti-CD3/CD28 antibodies plus FGL-1 FD WT (monomer), Y244A/E245A mutant, or mouse IgG1 isotype control (mIgG1) coated beads (n = 3). NF-κB::eGFP data are expressed as percent activation, normalized to the MFI of anti-CD3/CD28 stimulated LAG-3+ Jurkat cells MFI (100%). Statistical analysis was performed using one-way ANOVA, Statistical significance was performed using one or two-way ANOVA, followed by Tukey or Šídák’s post hoc tests, respectively. Error bars indicate standard deviation (SD). **** p < 0.0001, *** p < 0.001, ** p < 0.01, * p < 0.05, ns = not significant (p > 0.05).

**Supplementary Figure 4.**
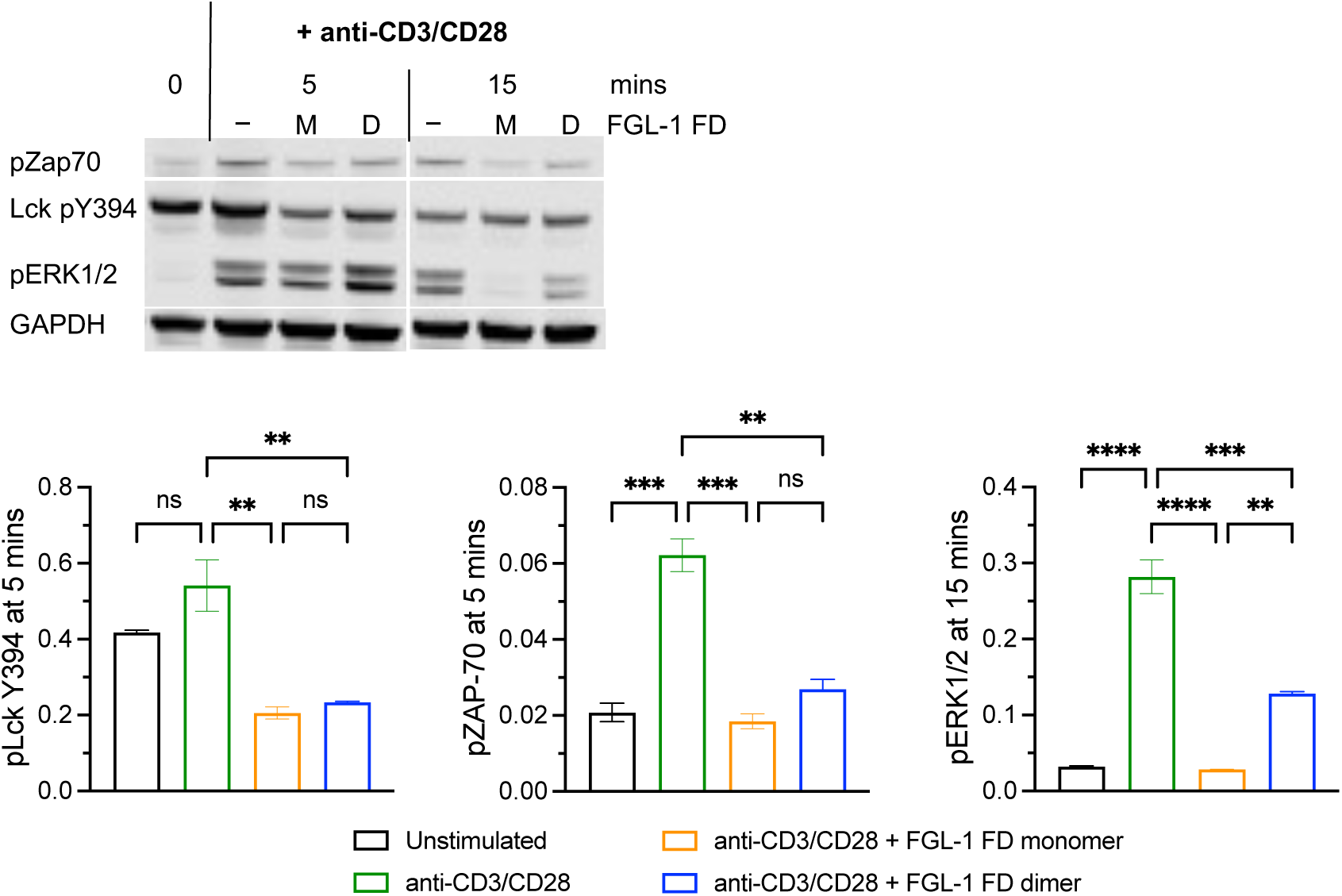
LAG-3/FGL-1 FD interaction inhibits TCR signaling within 5 minutes of stimulation. Western blot analysis of LAG-3+ Jurkat cells stimulated with anti-CD3/CD28-coated beads for 5-15 minutes, in the presence or absence of FGL-1 FD monomer or dimer (n = 3). Statistical analysis was performed using one-way ANOVA, followed by Šídák’s post hoc tests. Error bars represent standard deviation (SD). **** p < 0.0001, *** p < 0.001, ** p < 0.01, ns = not significant (p > 0.05).

**Supplementary Figure 5.**
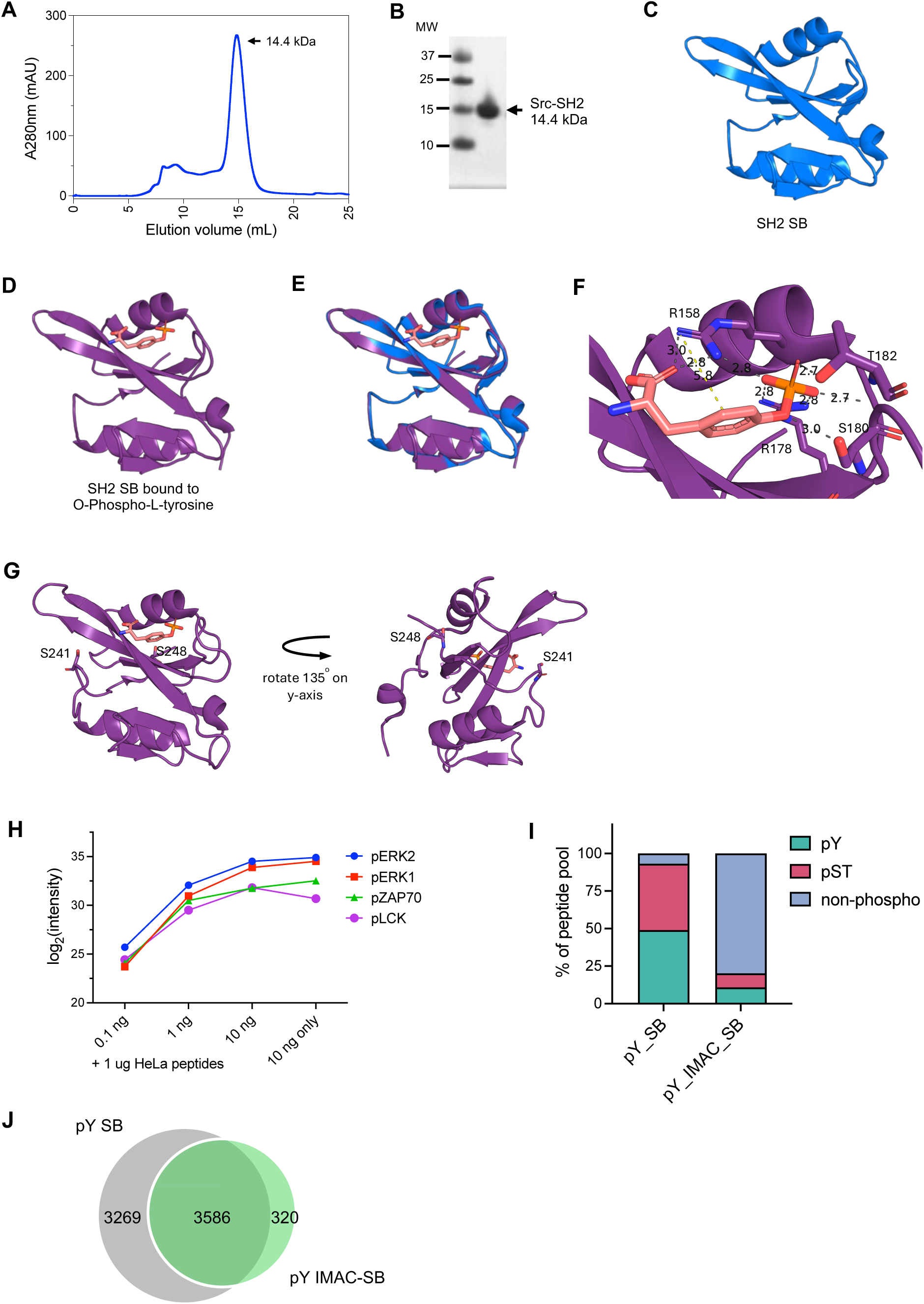
Purification and functional validation of recombinant Src SH2 superbinder (SB) domains. (**A**) Size exclusion chromatography elution profile of recombinant Src SH2 SB on a Superdex 75 Increase 5/150 GL column. **(B)** SDS-PAGE analysis of purified Src SH2 SB from panel (**A**) confirming high purity. Arrows indicates bands at the expected molecular weight. **(C, D)** High-resolution crystal structures of Src SH2 SB: apo form displayed in blue (**C**) and O-phospho-L-tyrosine (pY)-bound form in purple (**D**), both resolved at 1.5 Å. (**E**) Structural alignment of apo and pY-bound Src SH2 SB structures. Root Mean Square Deviation (RMSD) = 0.137 Å, indicating high structural similarity. (**F**) Detailed view of the pY binding pocket. Interacting side chains of Src SH2 SB residues (Arg158, Arg178, Ser180, Thr182) are shown as stick models; backbone ribbon in grey. Hydrogen bonds and salt bridges are represented with grey dashed lines; π–π stacking interactions with yellow dashed lines. (**G**) Structural visualization of Cys241Ser and Cys248Ser mutations as stick models. These residues are located distal to the pY binding site, confirming that the mutations do not affect pY-binding or structural integrity. (**H**) Quantitative mass spectrometry analysis of Lck pY505, ZAP-70 pY319, ERK1 pY204, and ERK2 pY187 phosphopeptides following enrichment with Src SH2 SB. Peptide intensities correlate with input concentrations, validating functionality. (**I**) Comparison of phosphopeptide enrichment methods using Src SH2 SB alone (pY SB) versus Ti^4+^ –IMAC plus SH2 SB (pY IMAC-SB) in H_2_O_2_-treated Jurkat T cell lysates (2mg). (**J**) Venn diagram showing overlap of pY peptides identified by pY SB and pY IMAC-SB workflows, highlighting shared and method-specific phosphopeptides.

**Supplementary Figure 6.**
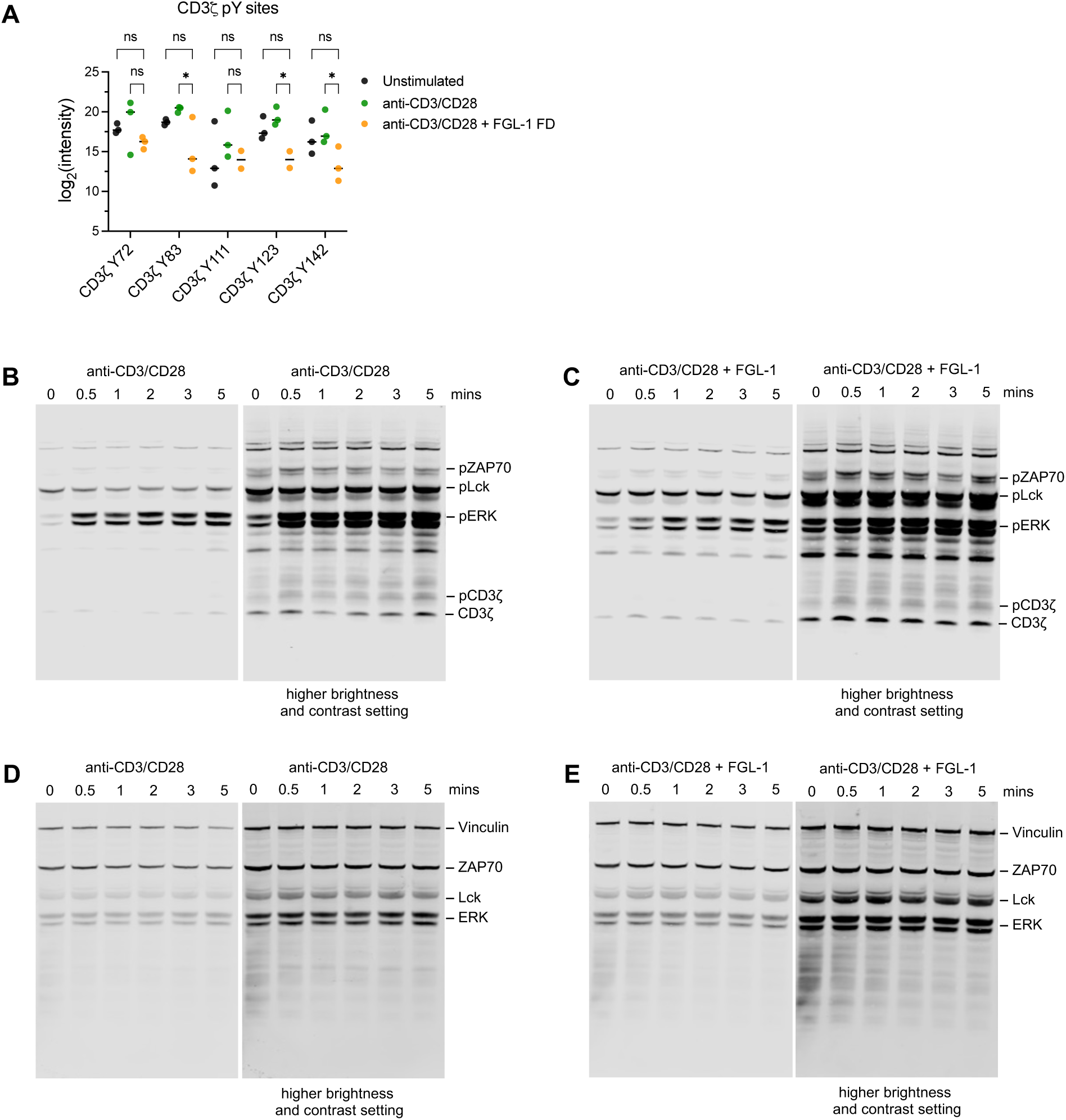
LAG-3/FGL-1 interaction inhibits proximal TCR signaling. (**A**) Peptide intensities of CD3ζ phosphotyrosine (pY) sites in unstimulated and anti-CD3/CD28-stimulated LAG-3+ Jurkat cells +/− FGL-1 FD, following 5-minute stimulation. Statistical analysis was performed using two-way ANOVA with Dunnett’s post hoc tests. * p < 0.05, ns = not significant (p > 0.05). **(B – E)** Representative western blot analysis of LAG-3+ Jurkat cells either unstimulated or stimulated with anti-CD3/CD28 +/− FGL-1 FD, probed for: **(B – C)** Phosphorylated proteins: CD3ζ Y142, ZAP70 Y319, and ERK 1/2 Y204/Y187 **(D – E)** Corresponding total protein levels., In each panel, the right-side blot displays a higher brightness and contrast setting to enhance visualization of low-intensity bands.

**Supplementary Figure 7.**
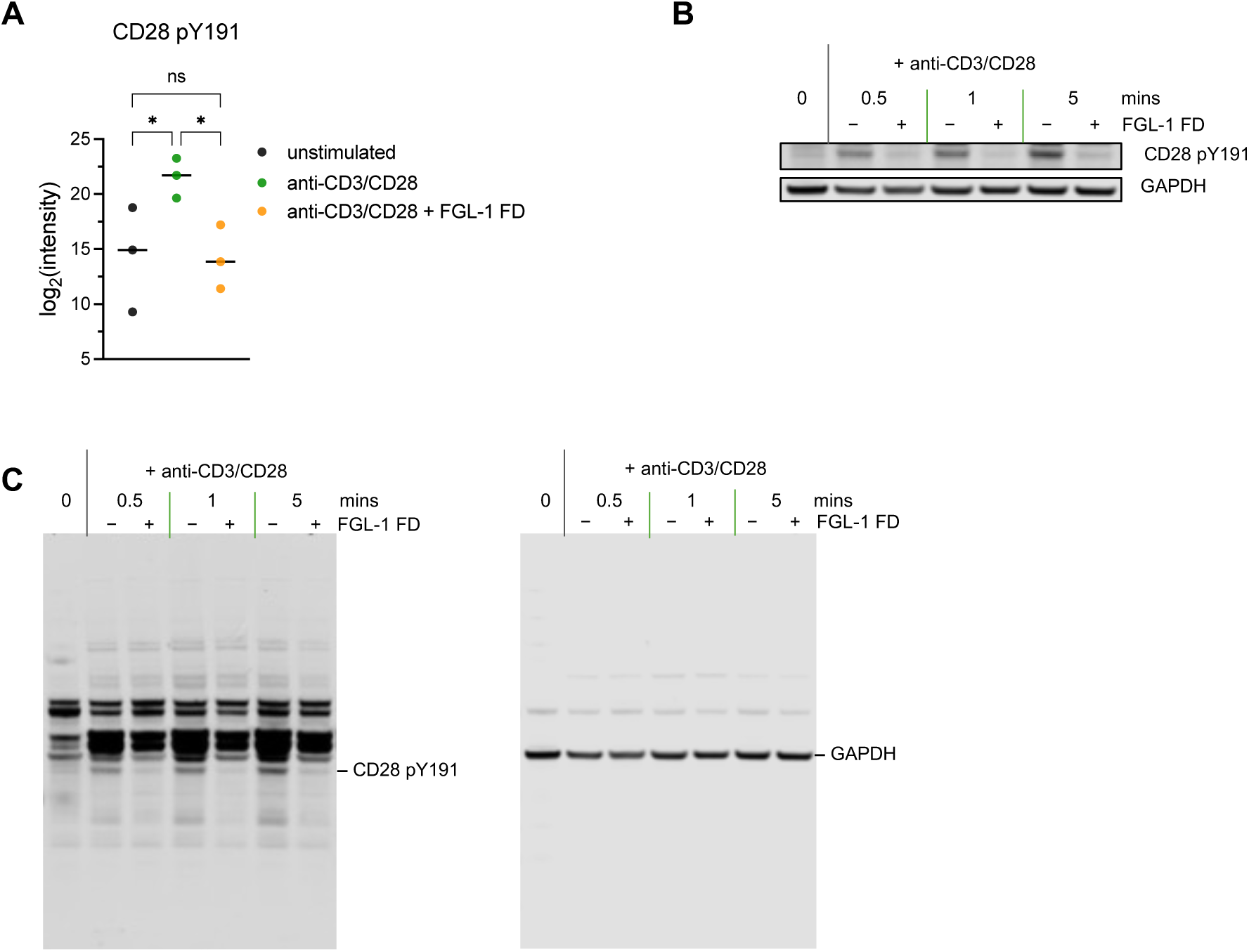
LAG-3/FGL-1 interaction suppresses CD28 Y191 phosphorylation. (**A**) Peptide intensities of CD28 phosphotyrosine Y191 in unstimulated and anti-CD3/CD28-stimulated with LAG-3+ Jurkat cells +/− FGL-1 FD, following 5-minute stimulation. Statistical analysis was performed using one-way ANOVA with Tukey’s post hoc tests. * p < 0.05, ns = not significant (p > 0.05). **(B – C)** Representative western blot analysis of CD28 Y191 phosphorylation in LAG-3+ Jurkat cells under the same stimulation conditions. **(B)** shows a cropped view highlighting the phosphorylated band, while **(C)** presents the full-length blot.

**Supplementary Figure 8.**
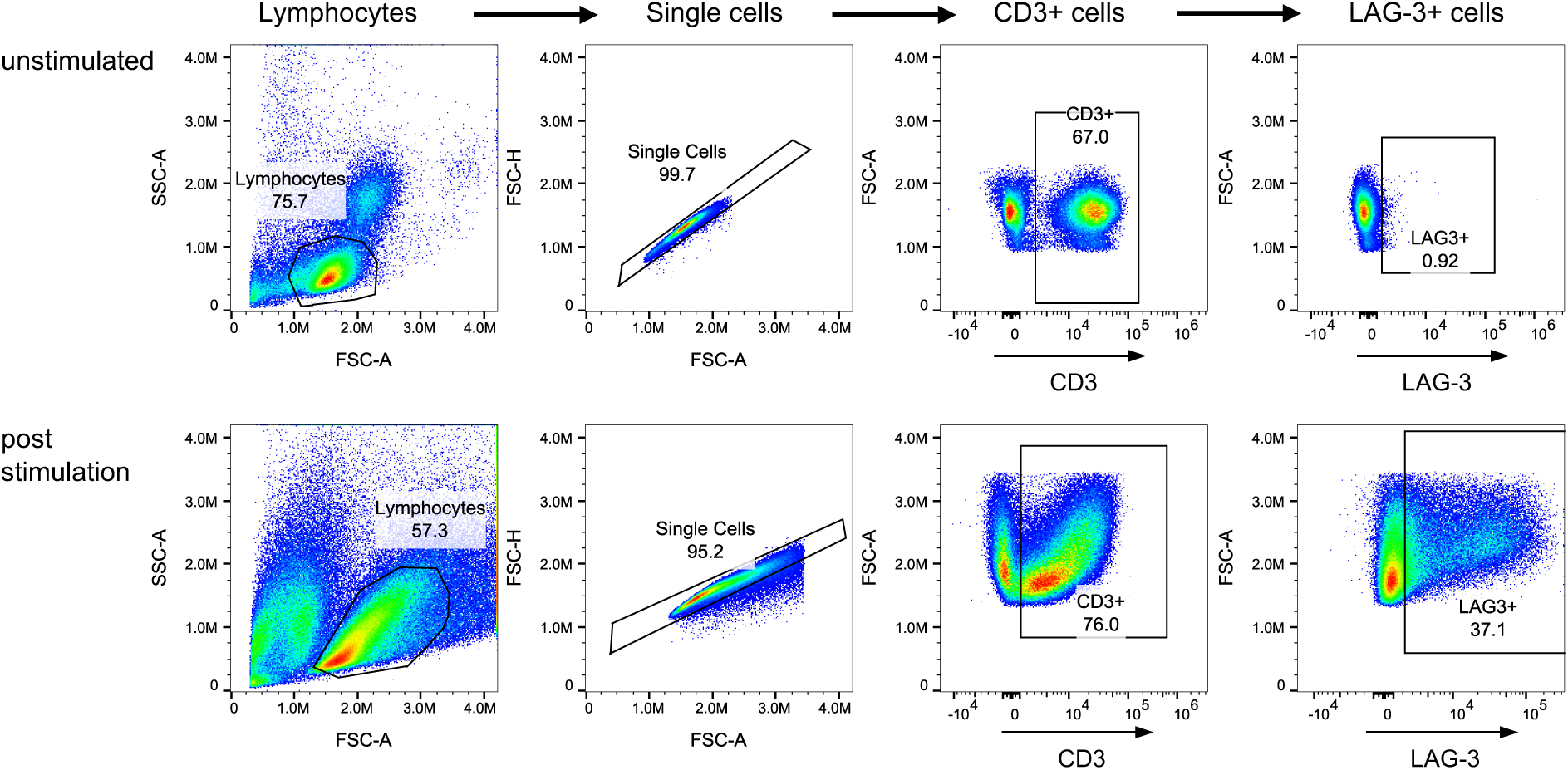
Flow cytometry gating strategy for primary T cells from healthy donors. Flow cytometry gating strategy used to identify CD3+/LAG-3+ from unstimulated and stimulated primary T cells derived from healthy donors, following stimulation to induce LAG-3 expression. CD3+/LAG-3+ cells were subsequently used for downstream analysis of phosphorylated protein expression, as shown in Figure 4.

**Supplementary Figure 9.**
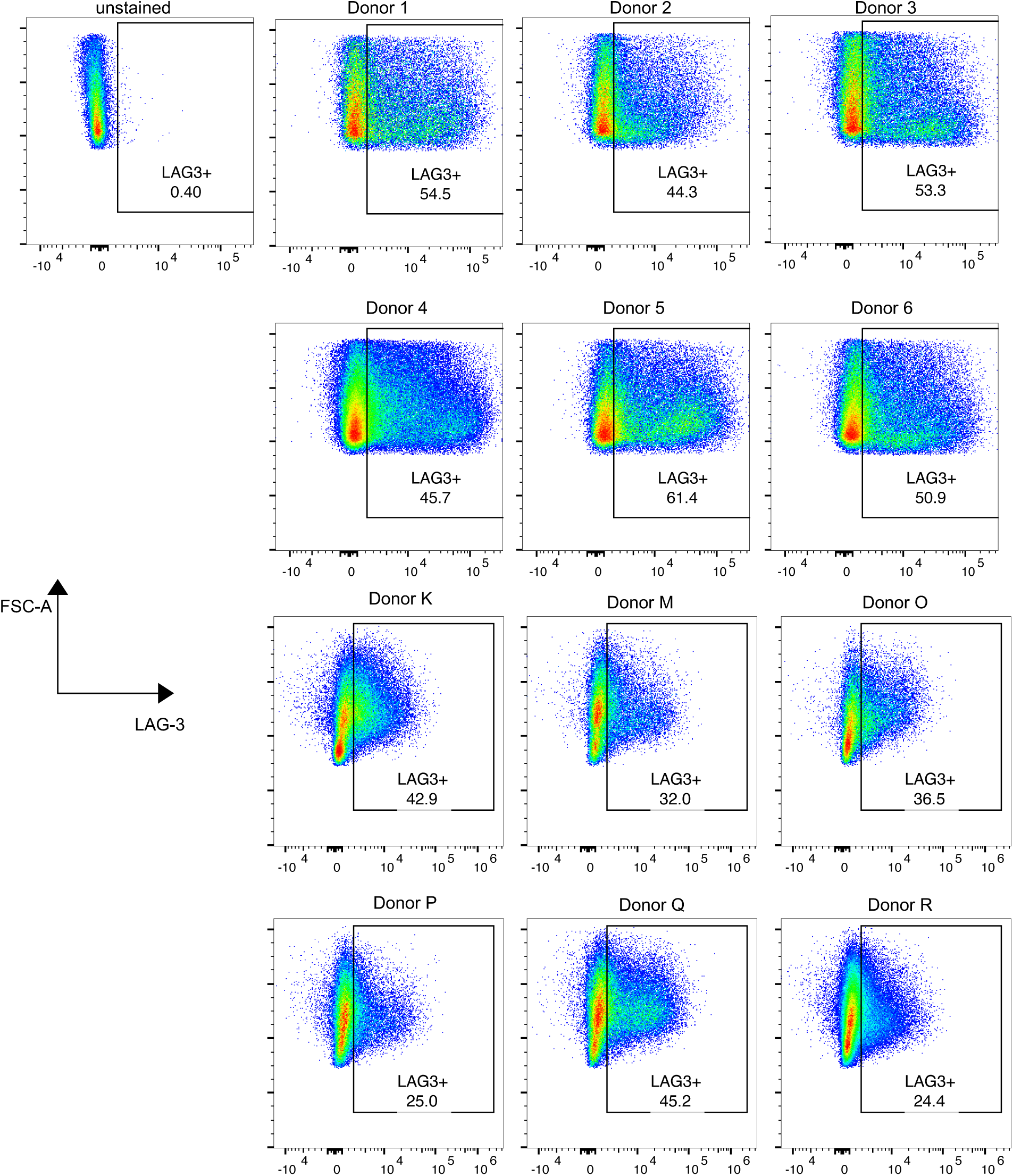
Induced LAG-3 expression on primary T cells. Flow cytometry plots showing LAG-3 expression in primary T cells from 12 healthy donor PBMCs following stimulation with anti-CD3/CD28 antibodies for 48 hours, followed by a 48-hour rest in IL-2.

**Supplementary Figure 10.**
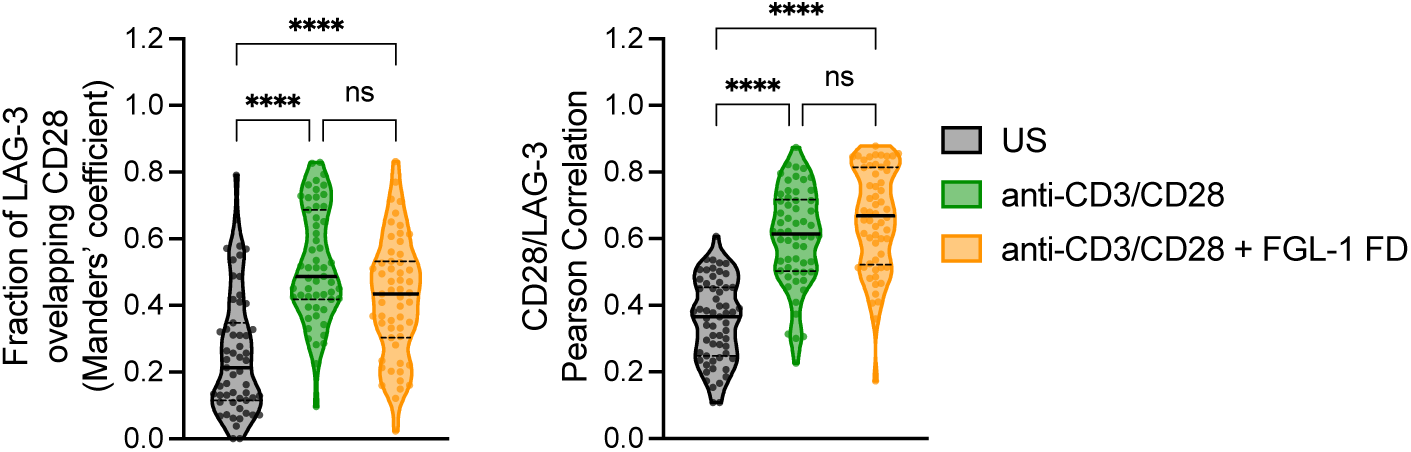
CD28 and LAG-3 colocalization analysis in unstimulated and stimulated +/− FGL-1 FD. MOC and PCC indicate an increase in CD28 and LAG-3 colocalization upon stimulation, with no significant difference between cells stimulated with and without of FGL-1 FD (n ≥ 60). Median values are shown as solid black lines, interquartile ranges as dashed black lines. Statistical significance was assessed using Kruskal-Wallis ANOVA, followed by Dunnett’s post hoc tests. **** p < 0.0001, ns = not significant (p > 0.05).

**Supplementary Figure 11.**
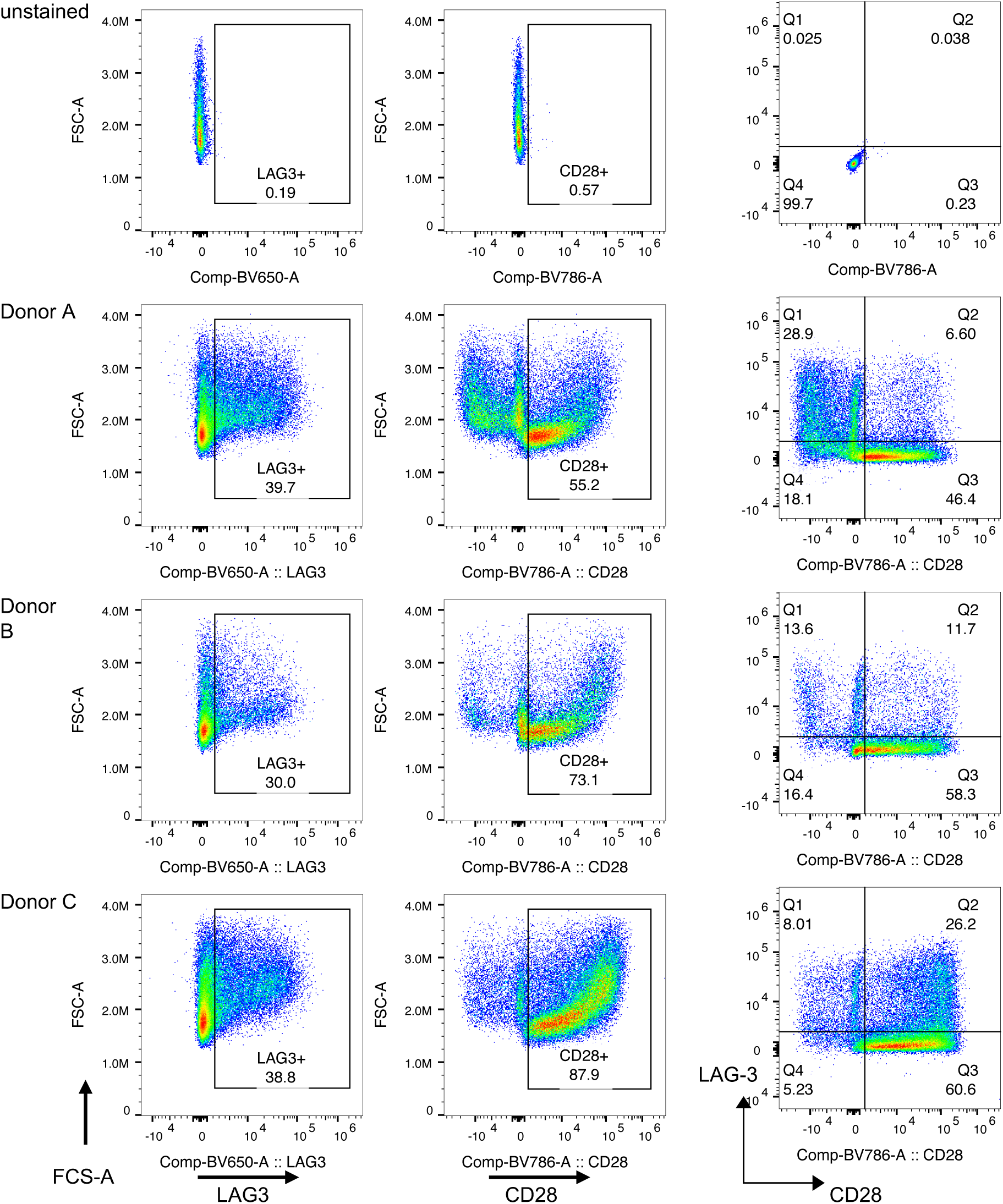
Variability in CD28 and LAG-3 expression in primary T cells. Flow cytometry plots showing CD28 and LAG-3 expression in primary T cells from four health donor PBMCs, following stimulation with anti-CD3/CD28 antibodies for 48 hours, and a 48-hour rest in IL-2. The gating strategy from **Supplementary** Figure 8 was used to identify LAG-3+ and CD28+ T cell populations.

## Supplementary Tables

**Supplementary Table 1.** FGL-1 FD Dimer X-ray diffraction data and refinement statistics.

|  | **FGL-1 FD Dimer** |
| --- | --- |
| **Space group** | **P2<sub>1</sub>22<sub>1</sub>** |
| Ave. unit cell | 39.78, 65.77, 90.01 |
| Resolution range (Å) | 37.2 – 1.74 (1.78 – 1.74) |
| Mean I/$\sigma$(I) | 9.6 (2.1) |
| Completeness of data (%) | 97.7 (78.7) |
| R<sub>merge</sub> | 0.09 |
| R/R<sub>free</sub> | 0.24/0.29 |

**Supplementary Table 2.** Src-SH2 SB X-ray diffraction data and refinement statistics.

**Supplementary Table. Src-SH2 SB X-ray diffraction data and refinement statistics**
| **PDB ID** | **8VCF** | **8VCG** |
| --- | --- | --- |
| Protein | Src-SH2 SB | Src-SH SB – pY |
| Space group | P6<sub>1</sub> | P6<sub>1</sub> |
| Ave. unit cell (Å) | 67.48, 67.48, 46.89 | 67.67, 67.67, 46.76 |
| Resolution range (Å) | 46.89 – 1.5 (1.53 – 1.5) | 29.30 – 1.61 (1.66 – 1.61) |
| Mean I/$\sigma$(I) | 22.5 (3.8) | 16.5 (2.1) |
| Completeness of data (%) | 100 (99.8) | 99.4 (95.3) |
| R<sub>merge</sub> | 0.07 | 0.07 |
| R/R<sub>free</sub> | 0.14/0.18 | 0.12/0.18 |

**Supplementary Table 3.**
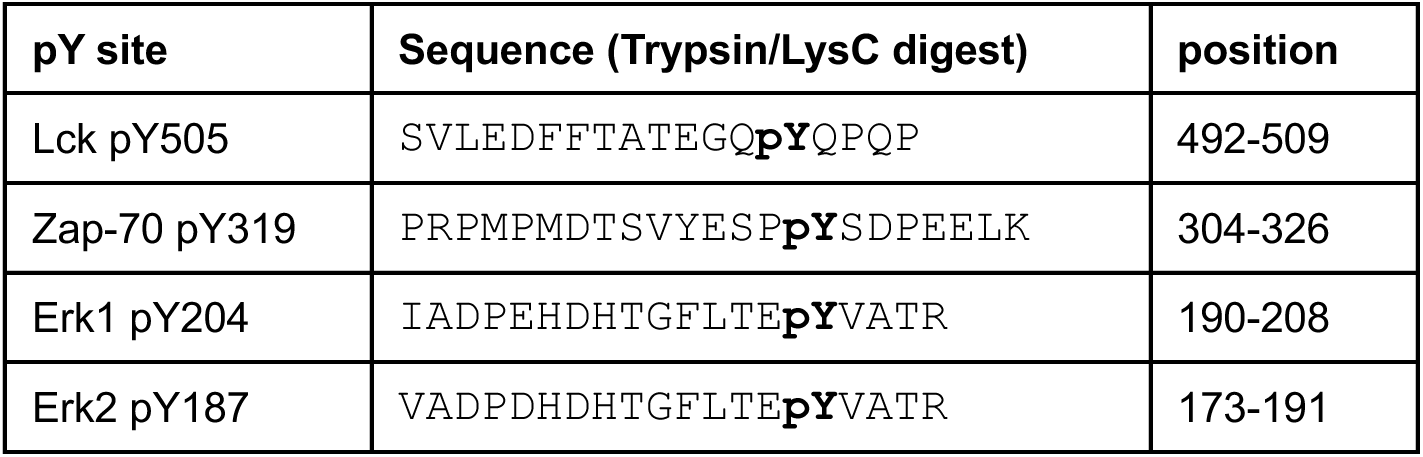
Peptide sequences of TCR signaling pY peptides.

**Supplementary Table 4.** Summary description of peptides identified using Src SH2 SB only and pY IMAC-SB enrichment methods.

| Sample | Total | Non-phospho | pS/T | pY | % pY |
| --- | --- | --- | --- | --- | --- |
| pY SB | 14017 | 965 | 6197 | 6855 | 48.9 |
| pY IMAC SB | 35870 | 28653 | 3311 | 3906 | 10.9 |

